# Conduction Velocity, G-ratio, and Extracellular Water as Microstructural Characteristics of Autism Spectrum Disorder

**DOI:** 10.1101/2023.07.23.550166

**Authors:** Benjamin T. Newman, Zachary Jacokes, Siva Venkadesh, Sara J. Webb, Natalia M. Kleinhans, James C. McPartland, T. Jason Druzgal, Kevin A. Pelphrey, John Darrell Van Horn, the GENDAAR Research Consortium

**Affiliations:** Department of Psychology, University of Virginia, Gilmer Hall, Charlottesville, VA 22903; UVA School of Medicine, University of Virginia, 560 Ray Hunt Drive, Charlottesville, VA 22903; School of Data Science, University of Virginia, Elson Building, Charlottesville, VA 22903; Department of Psychiatry and Behavioral Science, University of Washington, Seattle WA USA 98195; Seattle Children’s Research Institute, 1920 Terry Ave, Building Cure-03, Seattle WA 98101; Department of Radiology, Integrated Brain Imaging Center, University of Washington, 1959 NE Pacific St Seattle, WA 98195; Yale Child Study Center, 230 South Frontage Road, New Haven, CT 06520; Yale Center for Brain and Mind Health, 40 Temple Street, Suite 6A, New Haven, CT, 06520

**Keywords:** Conduction velocity, diffusion MRI, autism spectrum disorder, sex differences

## Abstract

The neuronal differences contributing to the etiology of autism spectrum disorder (ASD) are still not well defined. Previous studies have suggested that myelin and axons are disrupted during development in ASD. By combining structural and diffusion MRI techniques, myelin and axons can be assessed using extracellular water, aggregate g-ratio, and a novel metric termed aggregate conduction velocity, which is related to the capacity of the axon to carry information. In this study, several innovative cellular microstructural methods, as measured from magnetic resonance imaging (MRI), are combined to characterize differences between ASD and typically developing adolescent participants in a large cohort. We first examine the relationship between each metric, including microstructural measurements of axonal and intracellular diffusion and the T1w/T2w ratio. We then demonstrate the sensitivity of these metrics by characterizing differences between ASD and neurotypical participants, finding widespread increases in extracellular water in the cortex and decreases in aggregate g-ratio and aggregate conduction velocity throughout the cortex, subcortex, and white matter skeleton. We finally provide evidence that these microstructural differences are associated with higher scores on the Social Communication Questionnaire (SCQ) a commonly used diagnostic tool to assess ASD. This study is the first to reveal that ASD involves MRI-measurable *in vivo* differences of myelin and axonal development with implications for neuronal and behavioral function. We also introduce a novel neuroimaging metric, aggregate conduction velocity, that is highly sensitive to these changes. We conclude that ASD may be characterized by otherwise intact structural connectivity but that functional connectivity may be attenuated by network properties affecting neural transmission speed. This effect may explain the putative reliance on local connectivity in contrast to more distal connectivity observed in ASD.

## Introduction

Autism spectrum disorder (ASD) represents a complex, multifaceted condition having a significant sex bias in both diagnosis rate and behavioral symptom profile[1]. This sex difference has driven considerable research interest in determining diagnostic and sex-specific associations with brain structure and function as measurable using magnetic resonance imaging (MRI)-based techniques.

*ASD Neuroimaging:* MRI techniques have long been a focus in ASD for potential diagnosis and increased understanding of brain patterns that might explain symptom severity and clinical variation. Reviews of neuroimaging literature going back as far as 2003 report consistently detectable mean increases in total brain, parietal, temporal, and cerebellar volumes compared to typically developing (TD) comparison subjects, though there were apparent differences in findings depending on the age cohort being studied and an almost total lack of female subjects[2,3]. Using a large subset of male subjects from the Autism Brain Imaging Data Exchange (ABIDE), a more recent study has described widespread increased cortical thickness that diverges from TD during childhood but rapidly decreases to levels equivalent to TD participants upon entering adulthood[4]. Another study supported this finding, used a distance-based feature selection paradigm to find differences within paired ASD/TD subjects to see differences in cortical thickness that may map to areas key to functional networks. However, these areas of significant difference were relatively sparse throughout the brain[5]. Meanwhile, longitudinal studies examining volume changes during adolescence in key subareas such as the hippocampus and amygdala have shown no difference between TD and ASD[6]. While there is ample evidence that differential neuroimaging features are present to differentiate ASD from TD, machine learning techniques have only achieved 80% ROC classification success from combined fMRI and structural data across the brain[7]. It has also been questioned if there are reliable sex differences in ASD or if observed sex-by-diagnosis interactions map to areas typically observed to show sex differences during development[8,9]. Studies on sex differences have suggested that ASD males and females may not exhibit sex-specific differences seen in typically developing individuals[10]; however, experimental parameters such as small sample sizes, lack of consistent experimental control factors, and a relative lack of female subjects have generally impacted many studies, making it difficult to gain insight on the neurobiological basis for ASD from an imaging perspective.

*Connectomic Differences in ASD:* Altered functional connectivity in the brain is a widely noted feature of ASD. However, functional connectivity alterations are not uniform across the brain and are associated with a brain-wide pattern of hypo- and hyperconnectivity depending on the active region being examined[11–14]. A mega-analysis of nearly 2,000 individuals between 5-58 years old found that hypoconnectivity of the sensory and attentional network and hyperconnectivity between cortical and subcortical systems were associated with clinically relevant autistic features such as social impairments, repetitive behavior, and sensory processing[11]. This pattern is potentially present in early childhood, as a recent study by Yoon et al. (2022) found lower network efficiencies and reduced longer-range anterior-posterior resting-state connectivity in 2-6 year-olds with ASD[13]. Results from functional connectivity studies led to the development of a cortical underconnectivity theory, which postulated that the observed differences resulted from a deficiency in the presence or function of longer-range distal connections and an overreliance on shorter-range or local connections[15–17]. While initially controversial[18], several studies have supplied a variety of functional evidence in agreement with this theory[12,19–22]. Structural data has also provided evidence supporting reduced long-distance connectivity, with the corpus callosum (key to long-distance inter-hemispheric connectivity), in autistic individuals having lower fiber counts and volumes than comparison TD participants[23]. While the functional evidence describes the consequences of altered brain connections, it is essential to develop microstructural evidence that may highlight neuronal processes that lead to altered function to evaluate the underconnectivity theory and provide insight into the neurobiological basis for ASD.

*Neuronal Microstructure:* If the underconnectivity theory of ASD is accurate, it is likely that observed functional connectivity changes have a basis in the cellular microstructure of the neurons. Diffusion models of cellular microstructure can provide detailed information on architectural changes in brain structure at the sub-voxel level and describe specific cellular milieu rather than gross anatomy. In this framework, lower directional diffusion, referred to as *anisotropic*, and greater adirectional diffusion, referred to as *isotropic*, may indicate deficits in axonal morphology that would cause reduced action potential speed or efficiency and drive reduced connectivity deficits. Depending on which axons and axonal areas are affected, this might impact particular systems or longer/shorter range connections, particularly since longer-range connections rely on larger and greater diameter axons to efficiently transmit information across the brain[24–26]. Previous diffusion microstructure experiments have demonstrated that ASD participants display different axonal anatomy compared to TD participants, such as structural differences in neuronal white matter (WM), with ASD being associated with lower fractional anisotropy (FA) and increased diffusivity[27]. Using NODDI, one study identified increased extracellular water and decreased neurite density in ASD throughout major tractography bundles in the brain[28,29]. Other work has applied constrained spherical deconvolution to examine differences in axonal volume between ASD and TD participants using fixel-based analysis. In this context, a ‘fixel’ refers to an individual fiber population within a voxel, derived from the WM-FODs calculated using constrained spherical deconvolution, and allows for fiber-specific metric calculation relating to intra-axonal volume[30–33]. This analysis found that participants with ASD had significantly lower fiber density cross-section, a volume and fiber bundle diameter modified measure of intra-axonal volume, compared to TD participants across several major WM bundles of interest throughout the brain[27].

*Neuroimaging-based Assessment of Neuronal Conduction Velocity:* Beyond axonal fiber density, it is also possible to describe the relationship between axons and the myelin sheath surrounding them, providing the hallmark of the structural and diffusion microenvironment. The g-ratio is a quantitative proportion between axon diameter and myelin thickness first described by Rushton[34]. Increased myelin improves the resistivity of the axonal cellular membrane between the Nodes of Ranvier and provides for increased speed and energetic efficiency conducting action potentials[35,36]. Both axonal diameter and myelin thickness affect neuronal conduction velocity and spatial, energetic, and conductive properties within the human nervous system, leading to an optimal g-ratio of around 0.6-0.7[34,37,38]. The g-ratio, thus, describes a fundamental linkage between neurons’ structure and function (via conduction velocity). Genetic[39,40], electrical[41–45], and functional[8,46–49] evidence suggests that neural activity is disrupted in ASD. Post-mortem histological studies also demonstrate that myelin and axons display abnormal morphology in ASD[50]. MRI-based measures of g-ratio, conduction velocity, and microstructural axonal properties are needed to establish *in vivo* measurements to give insights into the underpinnings of ASD.

Developing more sensitive models of microstructure and axonal properties may provide insight into structural and cellular processes in ASD. The g-ratio is an important measure of axonal integrity, and that accounts for axonal diameter and the diameter of the myelin sheath surrounding the axon[51,52]. Conduction velocity is an important metric that addresses the capacity of the axonal fiber to carry information, and this can be derived, in part, from the g-ratio[53]. The present study aims to establish novel means of calculating both these metrics, in aggregate form across all axonal fibers in a voxel, using a combination of diffusion and structural MRI and applying these metrics to an ASD cohort.

## Methods

### Participants

Two-hundred seventy-three (mean age = 154.3 months ±35.21 S.D., age range = 96-216 months; 133 female [49%]) subjects from Wave 1 of an NIH-sponsored Autism Centers for Excellence Network were included in this study. Informed consent was obtained from all subjects and subject parents in accordance with requirements from the Institutional Review Board at the University of Virginia. All subject data were de-identified in accordance with the Health Insurance Portability and Accountability Act of 1996 (Public Law 104-191, 104th U.S. Congress). Neuroimaging and other study data are available from the National Institute of Mental Health Data Archive (NDA; Collection: #2021).The study cohort included 148 individuals diagnosed with ASD (mean age = 150.8 months ±34.31 S.D., 70 female [47%]) and 124 neurotypical participants (mean age = 154.3 months ±35.21 S.D., 62 female [50%]).

### Social Communication Questionnaire (SCQ)

All participants were assessed using the SCQ, a parent-report questionnaire comparing behaviors characteristic of ASD between ages 4-5 and present[54]. Scores on the SCQ range from 0-40 with higher scores indicating the presence of more behaviors characteristic of ASD. The SCQ is based on the Autism Diagnostic Interview – Revised (ADI-R)[55] and has been validated with the ADI-R and with ASD diagnosis[54]. A cutoff score of 15 has been reported to indicate ASD[56,57].

### Image Acquisition

Diffusion, T1-weighted, and T2-weighted images were acquired from each subject. Diffusion images were acquired with an isotropic voxel size of 2x2x2mm, 64 non-colinear gradient directions at b=1000 s/mm^2^ , and 1 b=0, TR=7300ms, TE=74ms. T1-weighted MPRAGE images with a FOV of 176x256x256 and an isotropic voxel size of 1x1x1mm, TE=3.3; T2-weighted images were acquired with a FOV of 128x128x34 with a voxel size of 1.5x1.5x4mm, TE=35.

### Image Data Processing

A simple overview of the image processing workflow, showing which imaging metrics contributed to calculation of other imaging metrics, is presented in Fig. 1. As described in Newman et al.[58], diffusion images were denoised[59], corrected for Gibbs ringing artifacts[60], and corrected for inhomogeneity fields using FSL’s *topup* and *eddy* commands utilizing outlier detection and replacemnet[61–63], and the final preprocessed diffusion images were up-sampled to an isotropic voxel size of 1.3x1.3x1.3mm^3^ [64]. WM, GM, and CSF tissue response functions were generated using the *Dhollander* algorithm[65], and single-shell 3-tissue constrained spherical deconvolution was used to generate the WM fiber orientation distribution (FODs) and GM and CSF representations. 3-Tissue Constrained Spherical Deconvolution[66–69] was used to calculate the voxel-wise maps of the fraction of signal arising from each of 3 compartments: an intracellular anisotropic, intracellular isotropic, and extracellular isotropic freely diffusing water compartment by setting the sum of all FOD coefficients equal to unity. WM-FODs were then used to create a cohort-specific template with a subset of 40 individuals counterbalanced between sex and diagnosis[70]. All subject’s WM-FODs were registered to this template using an affine non-linear transform warp, and then the template was registered to a b-value matched template in stereotaxic MNI space[71,72].

**Figure 1.**
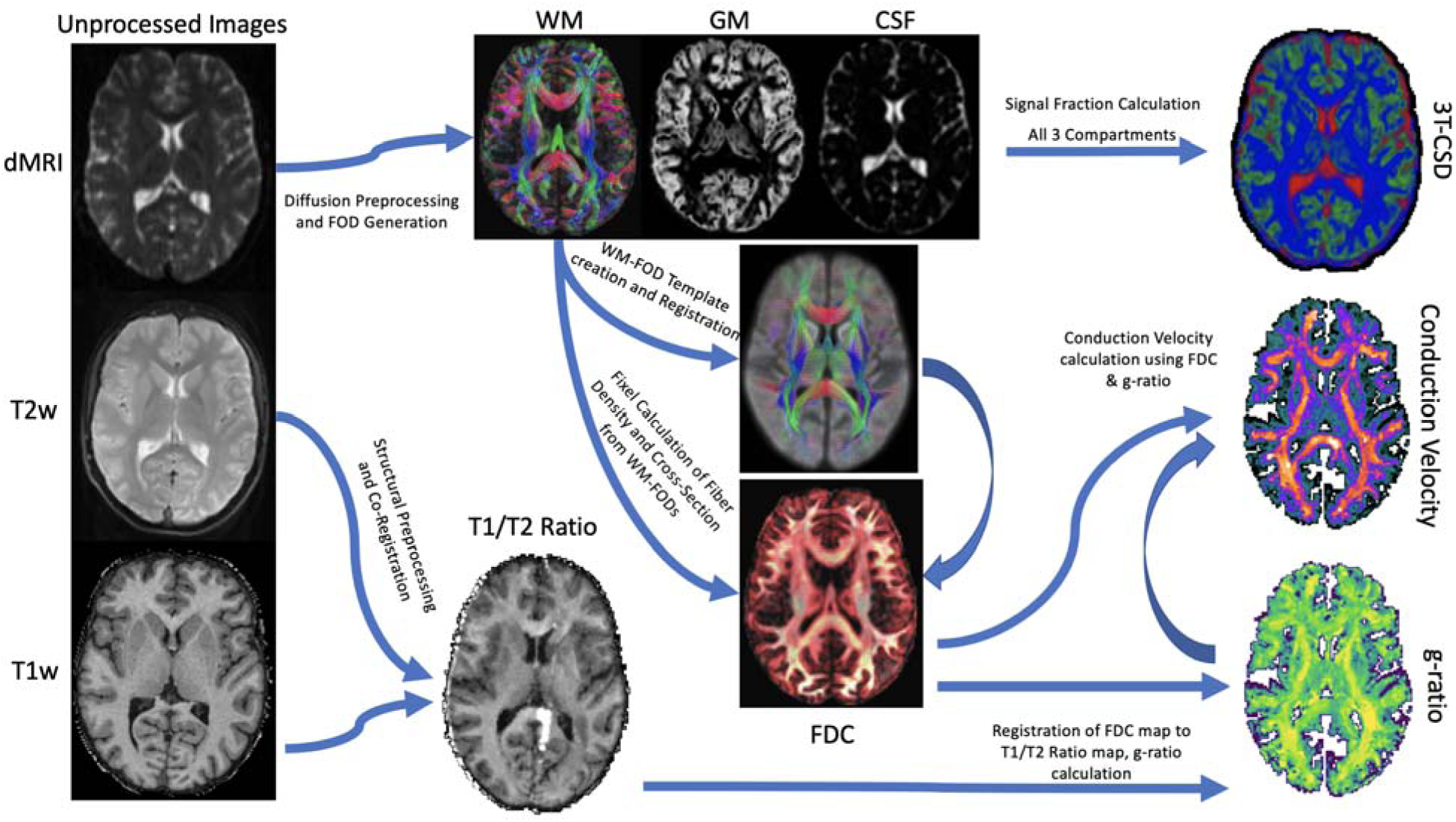
Simple flow diagram displaying which imaging metrics contributed to which output metrics. Fiber density cross-section (FDC) was derived from WM-FODs as part of a fixel analysis workflow, then summed voxel-wise as an intra-axonal volume fraction (AVF) estimate while T1W/T2W ratio was used as a myelin volume fraction (MVF) for the calculation of g-ratio and conduction velocity. It should also be noted that 3T-CSD metrics were separately registered to the MNI space atlases via a different procedure than the metrics derived from T1W/T2W ratio.

### Estimating Axonal Fiber Volume

A fixel-based morphometry (FBM)[30,31] approach was used to estimate the intra-axonal cross-sectional area within each voxel to be used as an apparent axonal volume fraction (AVF). FBM combines fiber density, computed from the integral of each WM-FOD spherical harmonic lobe, and cross-sectional area, computed using the Jacobian matrix determinant, which reflects the change in cross-sectional area when normalized to the template image. The cross-sectional change is used as a weight to the fiber density, and the two metrics are multiplied on a voxel-wise basis to obtain the fiber density cross-section related to the total intra-axonal volume (AVF)[30,33]. Each subject’s AVF maps were then registered to MNI space using the ANTs SyN nonlinear registration technique by aligning each to the 11-year-old adolescent template developed by Richards et al. Note a template approximately 1 standard deviation below the mean age of this study was used to better register the comparatively much smaller younger subjects[73,74].

### Myelination Density

T1w and T2w images were processed as described in the MICA-MNI workflow[75]; this involved performing N4 bias correction from ANTs[76,77] on both T1w and T2w images, rescaling both images from 0-100 based on their maximum values, co-registering the T2w to the T1w image using ANTs[77] rigid registration and then calculating the T1w/T2w ratio on a voxel-wise basis[78]. T1w/T2w ratio images were then registered to MNI space using the ANTs SyN nonlinear registration technique by aligning each to the same Richards et al. templates used to register the AVF maps. Despite recent findings that T1w/T2w ratio does not reliably quantify myelin in white matter regions[79], T1w/T2w ratio was selected as an estimate for myelin volume fraction for several reasons. The first was that more advanced means of quantifying myelin using specialized acquisitions such as magnetization transfer imaging or multi-echo T2 myelin water fraction imaging were not able to be implemented in this multi-site clinical consortium. Using more readily available T1-weighted and T2-weighted images allows our proposed methods for estimating aggregate g-ratio and aggregate conduction velocity to be applied across a far wider number of publicly available datasets. Finally, and perhaps most importantly from a microstructural standpoint, T1w/T2w ratio does reliably have increased values in cortical areas associated with greater myelination, such as the motor and sensory cortices[78,79]. This ease of use, potential for wide applicability, and sensitivity in cortical areas compelled the use of T1w/T2w ratio for myelin volume fraction estimation despite acknowledged shortcomings within white matter regions. There was no calibration or adjustments performed both because g-ratio values are generally not well established in adolescents and a desire to not alter or introduce additional error to g-ratio measurements before proceeding to aggregate conduction velocity calculation.

### Conduction Velocity Determination

The aggregate g-ratio was calculated on a voxel-wise basis according to Stikov et al. and was used according to Mohammadi & Callaghan as displayed in Equation 1[51,52,80,81]. As a measure of intra-axonal volume, the fiber density cross section was used as the AVF[31], and as a metric of myelin density, the T1w/T2w ratio was used as the myelin volume fraction (MVF). Both of these metrics represent the total sums of each respective compartment across the volume of the voxel and are a volume-based equivalent to the original formulation of g as the ratio of axon diameter (d) to fiber diameter (D).

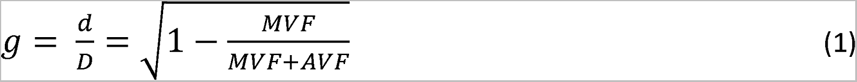

Aggregate conduction velocity was calculated based on the calculations of Rushton[34] and Berman et al.[53]; reiterating Rushton’s calculation that conduction velocity (θ) is proportional to the length of each fiber segment (l), and that this is roughly proportional to *D,* which in turn can be defined as the ratio between *d* and the g-ratio (g). Furthering the considerations of Rushton, Berman et al. show that a value proportional to conduction velocity can be calculated using axon diameter and the g-ratio as in equation 2[53]:

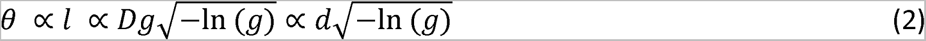

Herein, aggregate conduction velocity is calculated at the whole-voxel level, similar to Stikov et al.’s original conceptualization of an aggregate g-ratio, with each voxel representing an ensemble of g-ratios within each individual axonal segment with arbitrary orientation. Thus, instead of the direct measurement of axonal or myelin diameters, per se, the present study utilized a function of the arbitrarily oriented ensemble of intra-axonal volumes, returning to the AVF. It follows that increased or decreased diameter and axonal fiber density *in aggregate throughout a voxel* is proportional to intra-axonal volume. Additionally, these models were applied to voxels in both the white matter, where g-ratio has been calculated by several prior groups, to the cortex, which has been far less commonly examined in this way. As this was an adolescent cohort, a period that features widespread innervation of the cortex and development of myelin[82,83], we were especially interested in investigating fibers that entered the cortex. While it is more difficult to detect signal from axonal fibers in areas where fibers do not predominate, our use of SS3T-CSD is able to detect and remove signal from isotropic sources that may contaminate the axonal signal. Further, T1w/T2w ratio has been shown to more accurately track myelin content in cortex compared to deep white matter[79].

All imaging metrics, 3T-CSD signal fraction compartments, T1w/T2w ratio, aggregate g-ratio, and aggregate conduction velocity (Fig. 2) were averaged across each of 214 ROIs taken from the JHU-ICBM WM atlas (48 ROIs)[84] and the Destrieux Cortical Atlas (164 ROIs)[85]. Additionally, two composite ROIs were included, one of all 48 JHU ROIs and one of 150 neocortical regions from the Destrieux Atlas.

**Figure 2:**
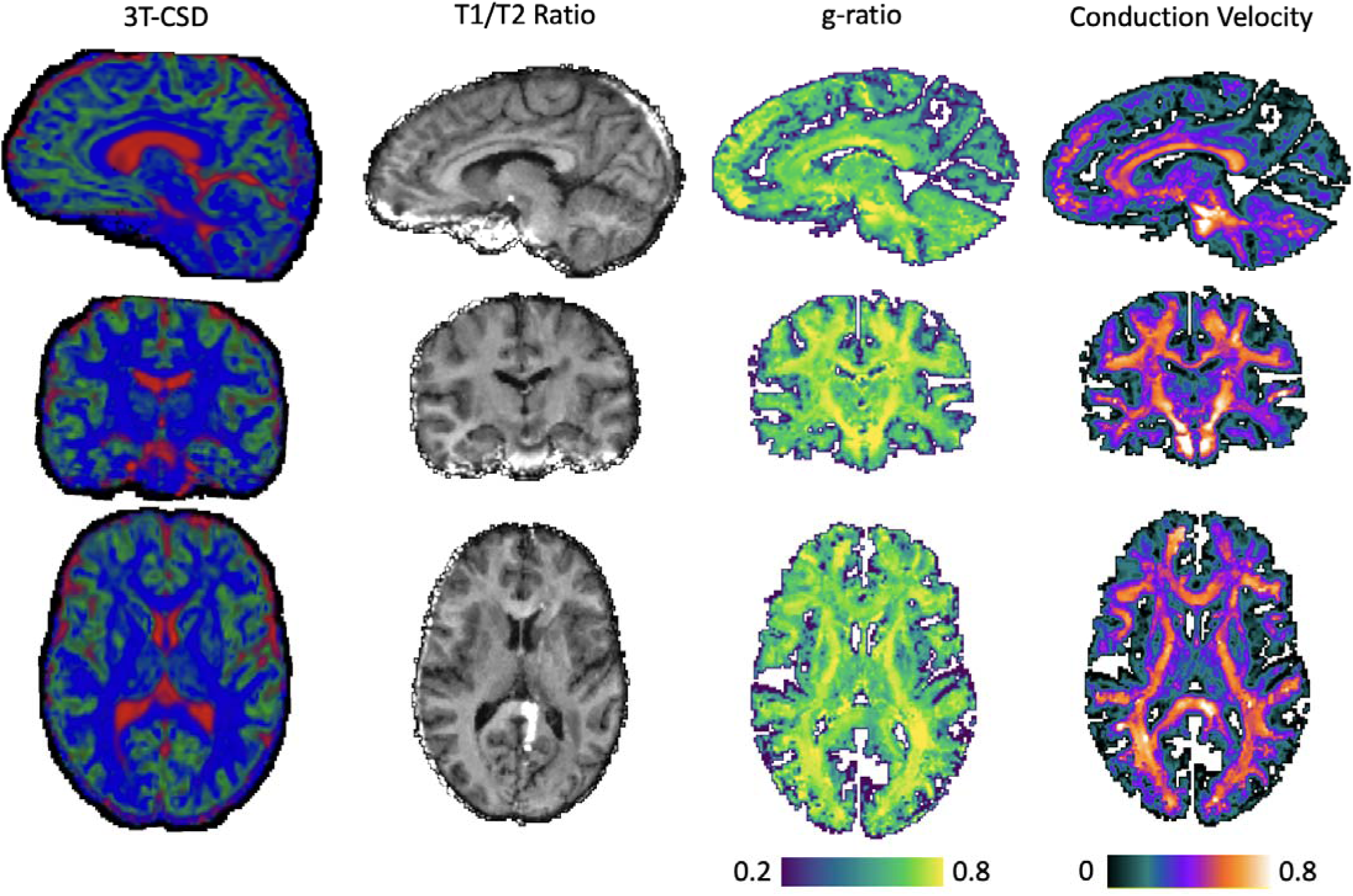
Illustration of each brain cellular microstructure metric from a single individual in this study. 3T-CSD is displayed as a composite of each of the 3 constituent tissue compartments having a value for each voxel, with blue voxels representing majority ICA (WM-like) signal, green voxels representing majority ICI (GM-like) signal, and red representing majority ECI (extracellular free water) signal. T1W/T2W ratio, aggregate g-ratio, and aggregate conduction velocity are colored with arbitrary scales to emphasize differences between brain areas.

### Statistical Approach

General linear models (GLMs) were used to test the relationship between each microstructural metric (intracellular anisotropic, intracellular isotropic, extracellular free water, T1w/T2w ratio, aggregate g-ratio, and aggregate conduction velocity) on diagnosis and sex differences as well as on SCQ total score with sex as a control. All GLMs featured subject age, IQ, scanner/site, and intracranial volume as control terms. All sets of microstructural results were corrected for multiple comparisons using the Benjamini & Hochberg (1995)[86] method across all 214 ROIs.

## Results

Given how the aggregate g-ratio and aggregate conduction velocity metrics were derived from diffusion and T1w/T2w ratio information, it is important to demonstrate that these metrics are not merely correlated with one another but actually describe different microenvironments in the brain. To test this, these metrics were first compared using linear models across all 214 ROIs with subsequent Benjamini and Hochberg correction for multiple comparisons. We aim to show that the metrics vary differently in two composite areas of the brain, the cortex and white matter. These results can be viewed in Fig. 3, and as expected, the methods derived from T1w/T2w ratio tended to be largely in agreement in both axonal ROIs from the JHU-ICBM atlas and in cortical ROIs from the Destrieux atlas. There were also many ROIs in the axonal areas where the ICA signal fraction was significantly associated with T1w/T2w ratio. This is especially notable as the WM-FODs that contribute to the ICA signal fraction are used to derive the fixel-based estimate of AVF. However, far less significant associations existed between ICA and either aggregate g-ratio or aggregate conduction velocity. In the cortex, despite the predominance of ICI signal, there was a widespread negative association between ECI (extracellular water) signal fraction and aggregate g-ratio and aggregate conduction velocity, suggesting that in this adolescent cohort, changes in cortical microstructure are primarily increases and decreases between intra-axonal and extracellular volumes, with no significant changes in ICI. We further performed two validation experiments to compare relative AVF values to known histological measurements in the corpus callosum and to observe the distribution of g-ratio values calculated across the brain of subjects, finding both in line with expectations from prior works (Fig. 4).

**Figure 3.**
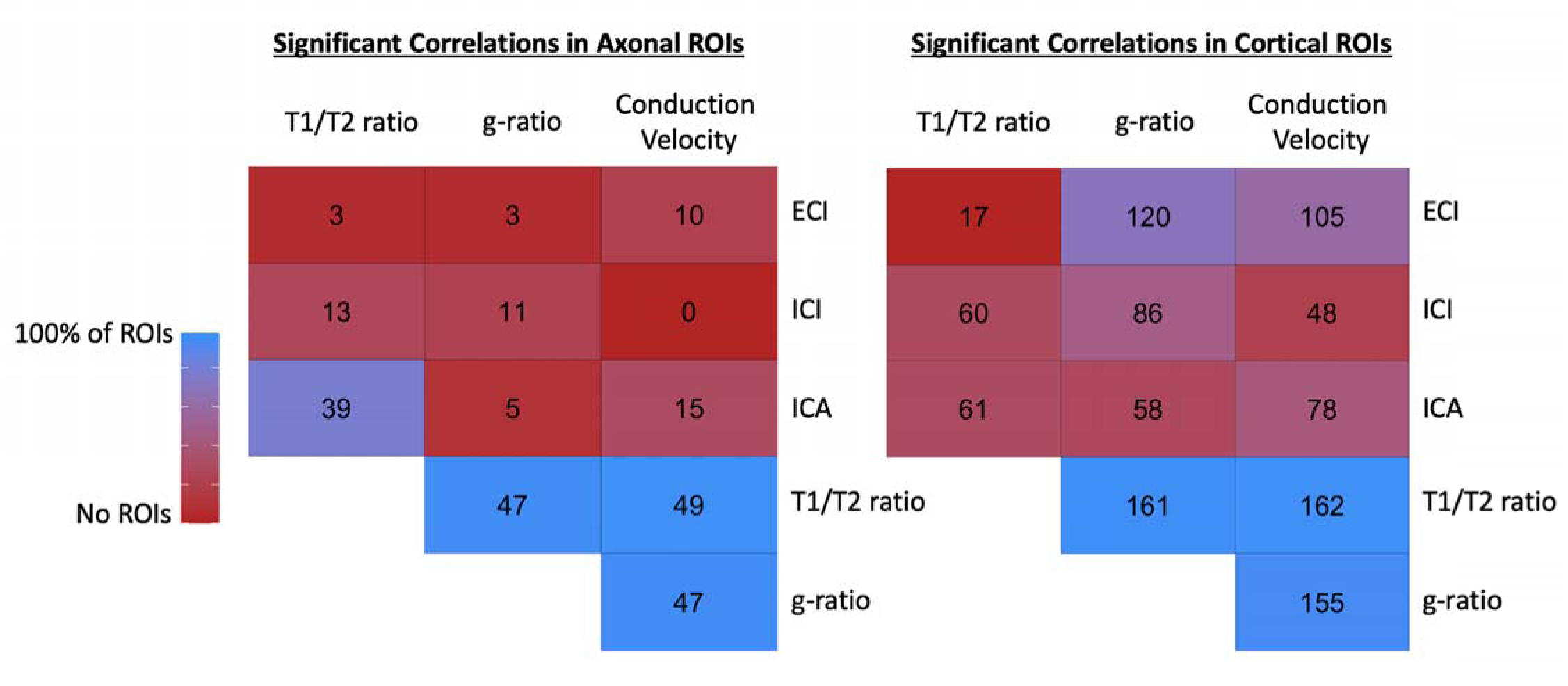
: Matrix displaying number of ROIs across axonal areas and cortex that remained significantly associated within the same ROI with each of the different metrics used in this study. Metrics derived from T1W/T2W ratio tended to have near universal significant associations between themselves (T1W/T2W ratio, aggregate g-ratio, and aggregate conduction velocity). There was not very widespread associations in the axonal ROIs between metrics derived only from diffusion and metrics derived from T1W/T2W ratio except for the ICA signal fraction (which is intended to measure intra-axonal signal fraction). ECI (extracellular water) signal fraction had negative associations while ICI and ICA had positive associations with T1W/T2W ratio derived metrics. This was particularly true in the cortical ROIs where ECI signal fraction was significantly associated with aggregate g-ratio and aggregate conduction velocity in a majority of ROIs.

**Figure 4.**
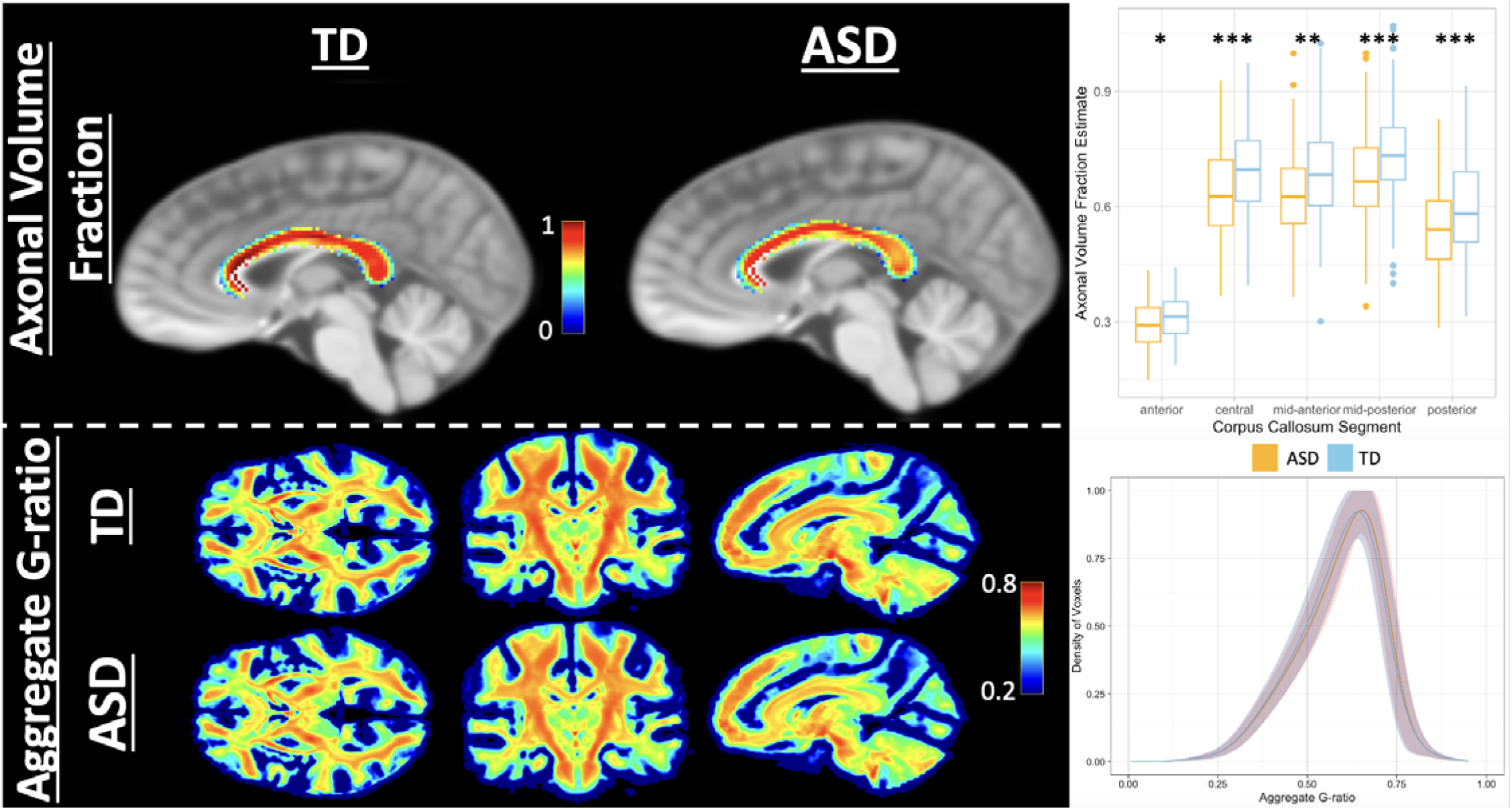
<u>: (Top)</u> Validation experiment examining mean AVF values derived from fixel-based analysis fiber density and cross-section in the corpus callosum of both ASD and TD subjects. Across both groups the AVF closely matched the pattern of axon diameter measured by histological means in humans by Aboitiz et al., (1994) with the lowest values in the anterior (genu) of the corpus callosum, higher values in the posterior (splenium), and the highest values in the central, mid-anterior, and mid-posterior portions of the body[103,104]. Additionally in each of the segments of the corpus callosum the AVF (FDC) was significantly higher in TD than in ASD after controlling for sex, age, intracranial volume, and IQ and performing a Benjamini and Hochberg multiple comparison correction. (Bottom) Distribution of mean aggregate g-ratio across whole brain from both TD and ASD cohorts. The density curve of voxels (mean density, shadow is range across all subjects in each cohort) is centered close to the ideal 0.6 described by Rushton with a longer curve to the distribution for lower than mean values.

These metrics were then assessed using the full model for relationships to variables of interest. For full summarized results of all adjusted p-values, see Table 1, for raw means from each metric in every ROI, see S1 Table, for all p-values before and after adjustment, see S2-S7 Tables for ECI, ICI, ICA, T1w/T2w ratio, aggregate g-ratio, and aggregate conduction velocity respectively. Following multiple comparison corrections, 93 ROIs were found to have significantly greater extracellular water signal fraction in individuals with ASD compared to TD participants (Fig.5), including in both composite ROIs covering the WM and neocortex. The significant ROIs were almost entirely located in the cortex, and no individual JHU-ICBM WM atlas ROIs, other than the entire composite ROI, survived multiple comparison corrections (Fig.6). 7 ROIs located in the subcortical gray matter also displayed significantly greater extracellular water signal in ASD compared to participants. These included the nucleus accumbens, hippocampus, and thalamus on the left exclusively, and bilateral caudate and putamen. Areas with the strongest relationship between extracellular water signal and ASD were the bilateral caudate nuclei and a bilateral series of pre- and post-central gyri in the parietal, superior temporal, and superior frontal lobes (Fig.5).

**Figure 5:**
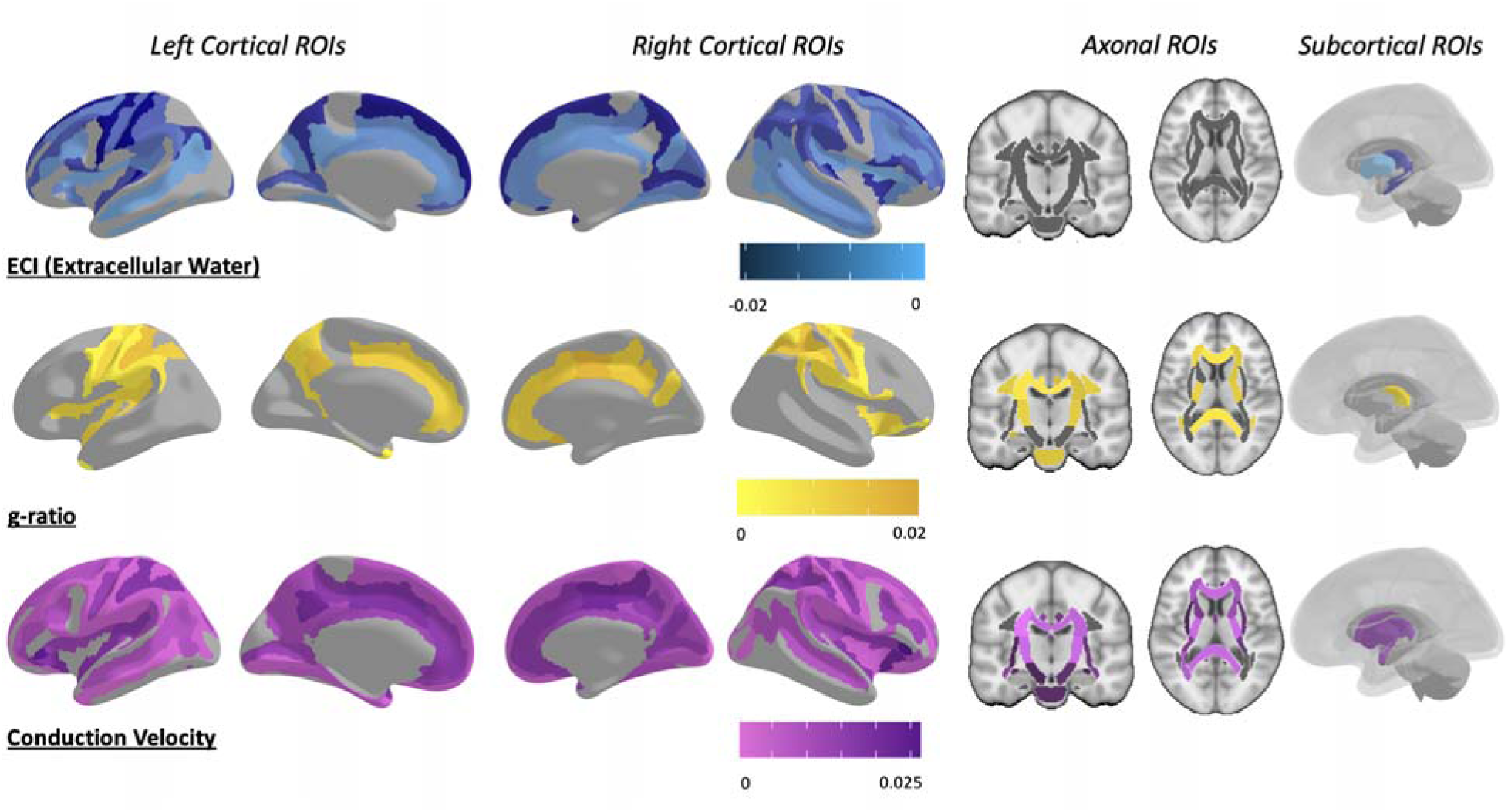
Map of ROIs that were significant at the p<0.05 level between ASD and TD after multiple comparison corrections. Colored by the slope of the relationship, with ASD individuals having greater extracellular water signal fraction, reduced aggregate g-ratio, and reduced aggregate conduction velocity in each of the significant ROIs. Aggregate conduction velocity was by far significantly different in more ROIs than any other metric, followed by the extracellular water compartment. However, there was a high degree of spatial specificity between the metrics, with aggregate g-ratio and aggregate conduction velocity involving extensive axonal areas and extracellular water differences primarily arising in the cortex and subcortex. See Supplementary Table 2, 6 & 7 for p-values for ECI, aggregate g-ratio, and aggregate conduction velocity, respectively.

**Figure 6:**
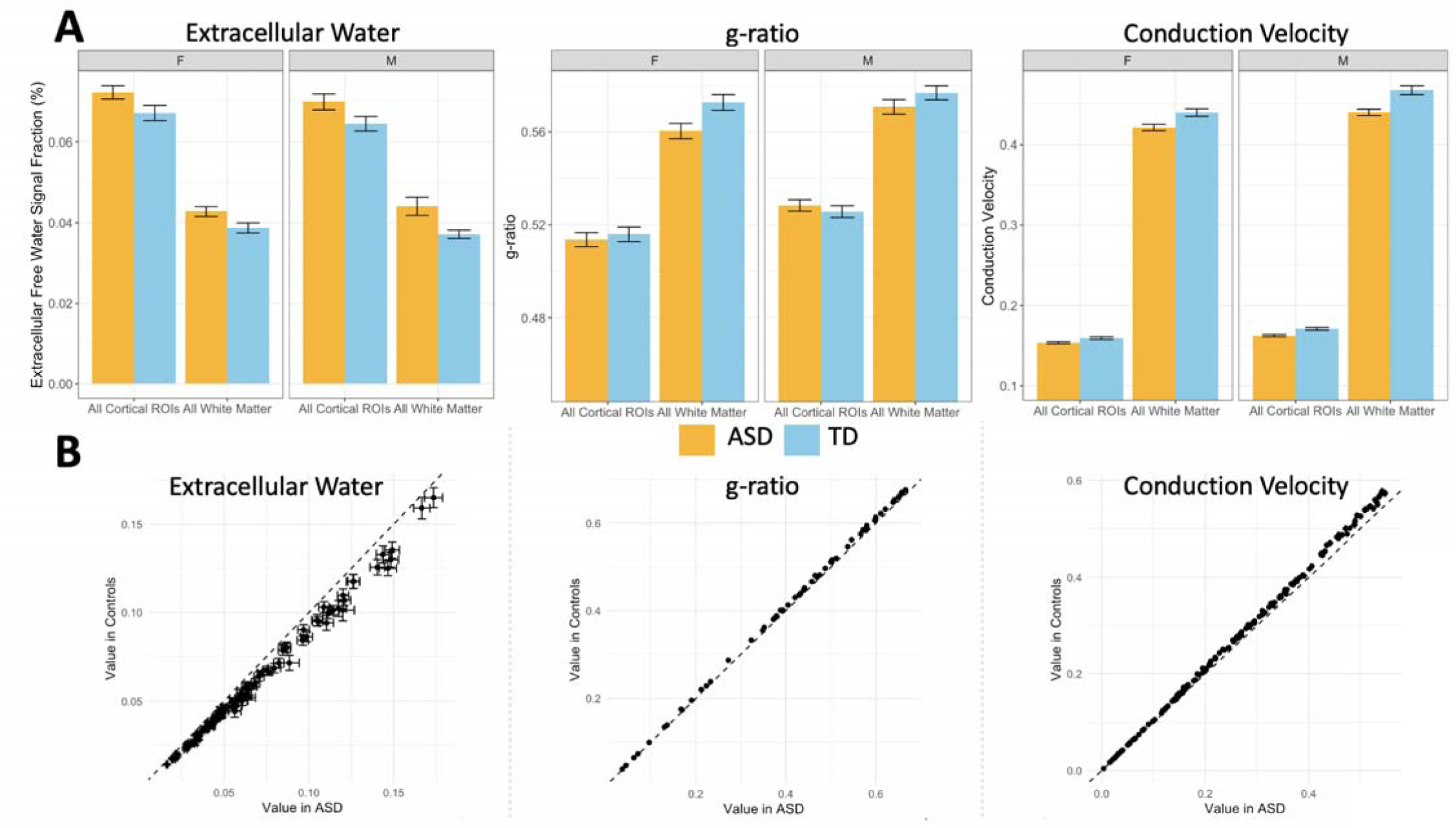
Chart displaying differences between extracellular free water, aggregate g-ratio, and aggregate conduction velocity levels in bilateral composite ROIs compared to TD participants, separated by sex. (A) Both boys and girls with ASD had significantly increased extracellular water, significantly decreased aggregate g-ratio, and significantly decreased conduction velocity. (B) Mean value in each significant ROI, after multiple comparison testing, with error bars representing standard error in each group. Dotted line represents x=y, showing that each individual ROI followed the directionality seen in the whole brain ROIs. Additionally, the effect appears to become more pronounced with higher mean value in extracellular value and aggregate conduction velocity.

**Table 1:**
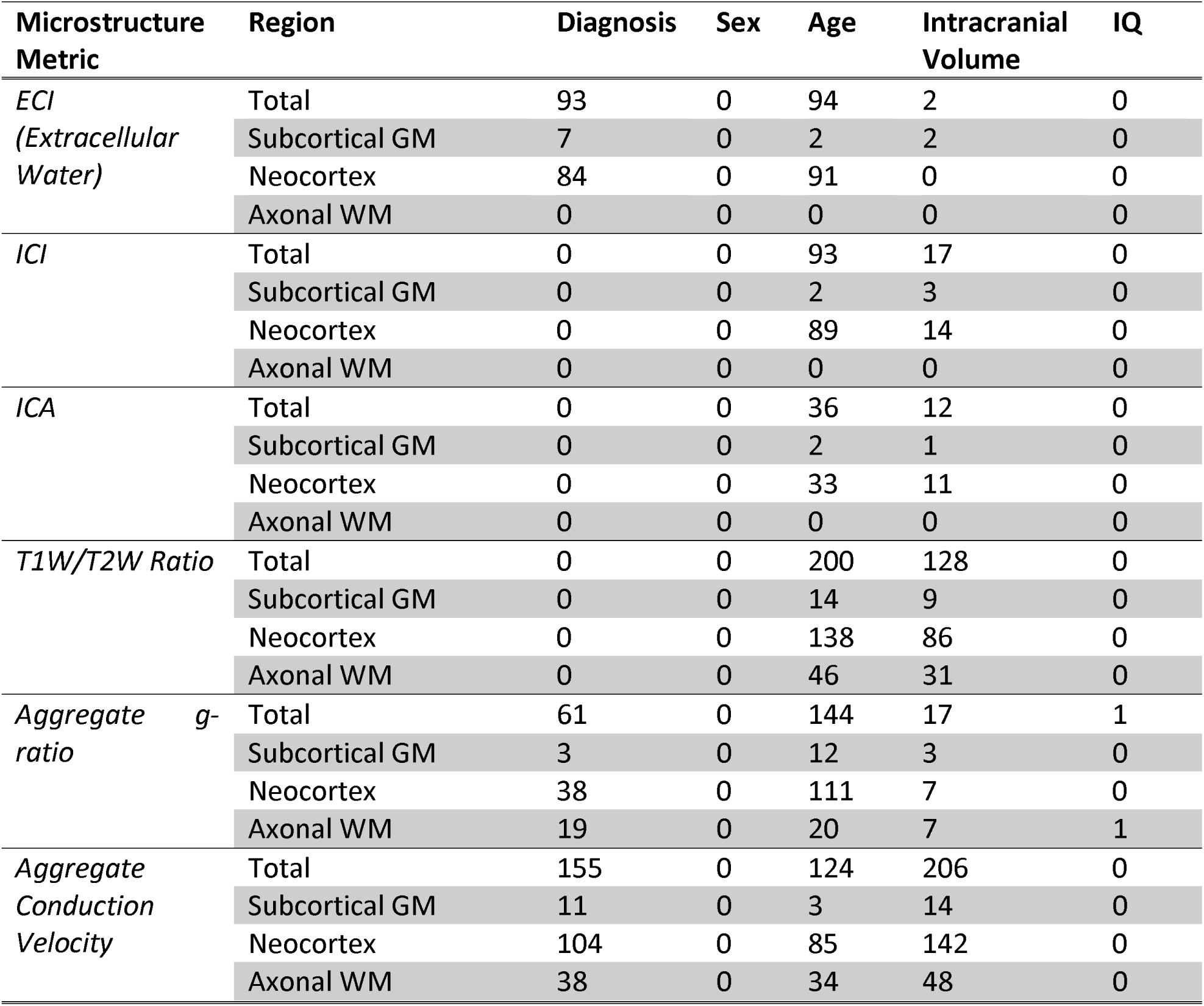
Table summarizing the number of ROIs from each microstructural metric and each anatomical subgroup that were significant at the p<0.05 level for each variable after multiple comparison corrections. The total value for each includes the 2 additional composite ROIs. The right cerebral peduncle was the only aggregate g-ratio significantly related to IQ.

Sixty-one ROIs were found to have significantly reduced aggregate g-ratio in ASD compared to TD participants (Fig.5), including the composite WM ROI but not the composite cortical ROI (Fig.6). Significant ROIs were located throughout the neocortex and axonal areas but were especially prominent in the superior parietal lobe. Only three subcortical areas showed significant differences, the right and left thalamus and the right nucleus accumbens.

One-hundred and fifty-five ROIs were found to have significantly reduced aggregate conduction velocity in ASD compared to TD participants (Fig.5), including both composite ROIs in the neocortex and WM (Fig.6). Significant ROIs included the majority of both atlases, with widespread associations between diagnosis and aggregate conduction velocity. Subcortically, there were significant associations with the bilateral amygdala, caudate, putamen, thalamus, and the right hippocampus. There was also a strong positive association between aggregate conduction velocity and intracranial volume, with 206 ROIs surviving multiple comparison corrections, including every WM ROI from the ICBM-JHU-WM atlas. Four of the non-significant ROIs were the bilateral temporal and occipital poles, and the remaining three were the left inferior occipital gyrus and sulcus, the right lunate sulcus, and the right lateral orbital sulcus. Receiver operator characteristic (ROC) analysis was performed using the ‘pROC’ package in RStudio on the linear model for the mean measurement made within each composite ROI to determine which microstructural metric with significant ROIs was most sensitive and specific in regards to subject diagnosis (Fig. 7)[87]. ROC analysis is a commonly used technique to test the performance of a diagnostic classification[88]. By assessing the sensitivity (proportion of true positives) and the specificity (proportion of true negatives) at multiple cutoff values along the range of observed data, the ability of a particular measure to separate two groups can be assessed. The area under the ROC curve (AUC) is a value that describes the overall accuracy of the particular classification metric and is used here to assess the effectiveness of each microstructural metric in each composite ROI to correctly determine a subject’s diagnosis. The ECI signal fraction within the composite WM ROI had the highest AUC (0.89), indicating the best sensitivity and specificity to diagnosis, while aggregate conduction velocity had AUC values in excess of 0.70 in both composite ROIs. The best performing composite ROIs from each microstructural metric are compared to age in Fig. 8, showing relatively consistent differences between TD and ASD across the cross-sectional age range of the study suggesting that observed differences are not related to differentially timed maturation or pubertal initiation.

**Figure 7:**
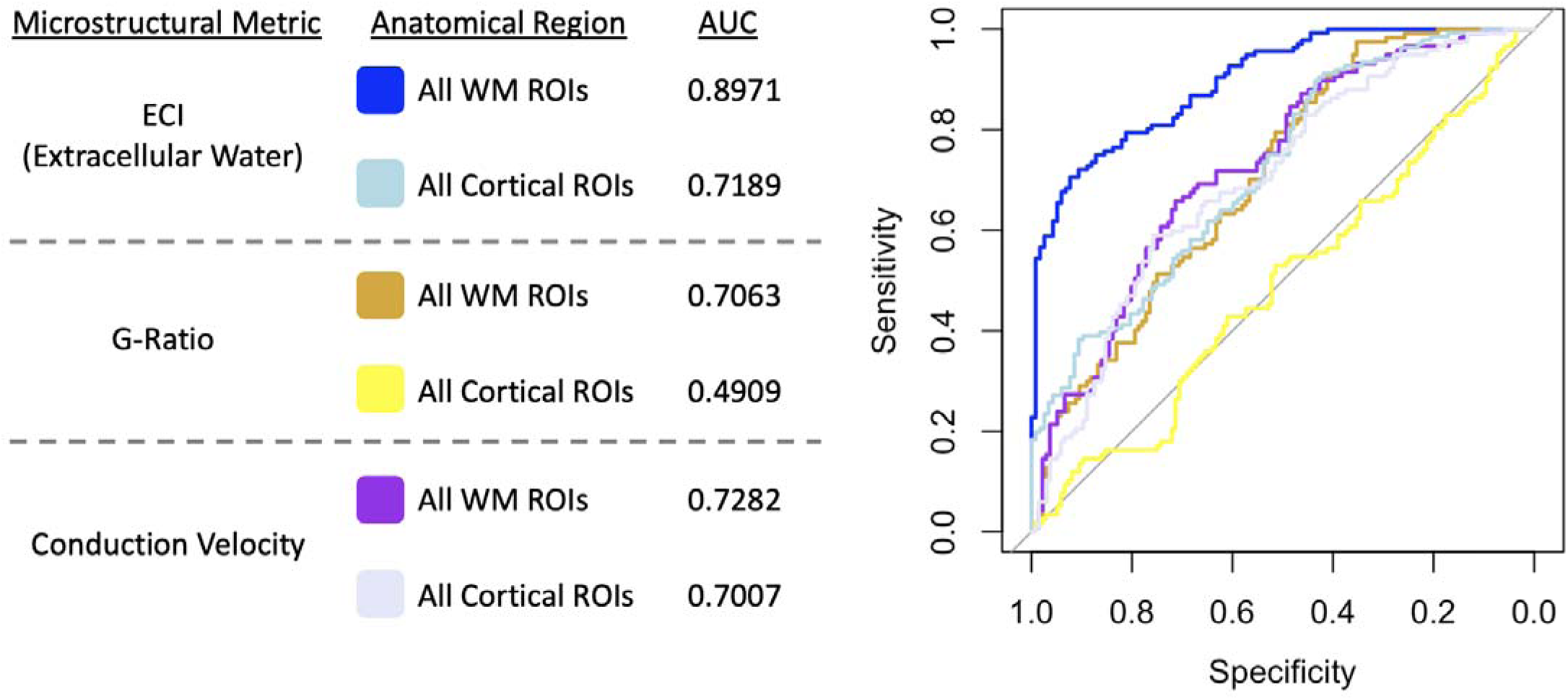
ROC analysis of the 3 microstructural metrics measured within the composite ROIs (axonal WM ROIs and Cortical ROIs, from the JHU WM atlas and Destrieux atlas) including AUC calculation. The same linear model as the ROI analysis was used to predict classification of subject diagnosis. Analysis indicated that ECI signal fraction within the composite WM ROI had the highest sensitivity and specificity to subject diagnosis, followed by aggregate conduction velocity within the composite WM ROI and ECI within the composite cortical ROI. Note that aggregate g-ratio had a sensitivity and specificity of approximately chance within the composite cortical ROI however this ROI was not statistically significant in the primary regional analysis.

**Figure 8.**
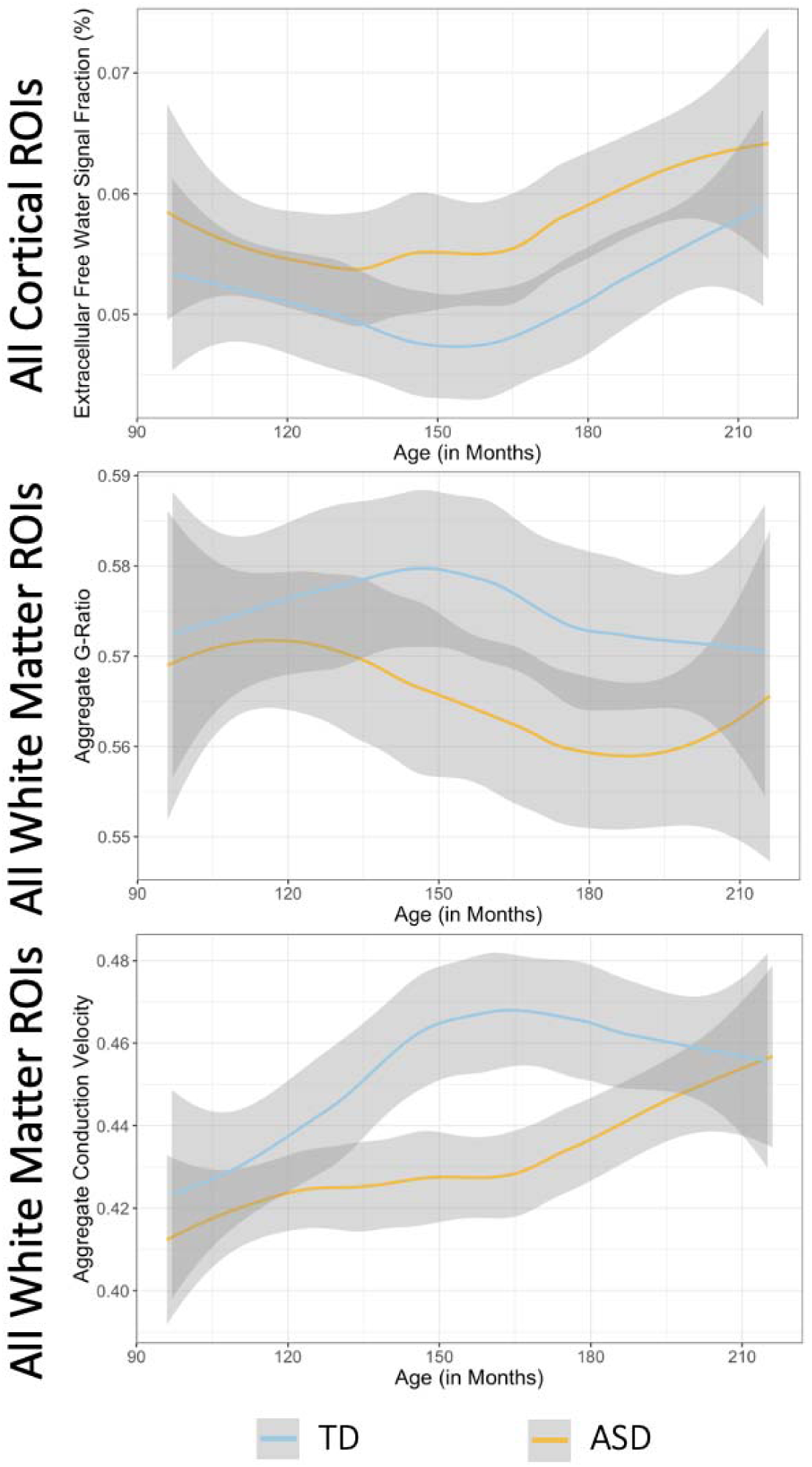
Chart displaying the relationship between subject age and the microstructural metrics significantly related to diagnosis (ECI signal fraction, aggregate g-ratio, and aggregate conduction velocity) within the top performing ROIs from the ROC analysis (Fig. 7). Curve is a loess line with a 0.95 confidence interval shadow. In each composite ROI differences between ASD and TD appear largely consistent across age, ECI and aggregate conduction velocity display positive relationships with age while aggregate g-ratio displays a negative relationship with age.

In regard to diagnosis, no other microstructure metric (neither ICI, ICA, nor T1w/T2w ratio) survived multiple comparison corrections in any ROI. There was no significant effect of sex or of sex-by-diagnosis interaction in any ROI after multiple comparison corrections (Fig. 9). Though there were no significant associations with sex in any microstructural metric in any ROI, diagnosis groups show varying relationships with age that are not consistent across sexes, with far larger T1w/T2w ratio values in TD males compared to TD females, while in ASD, females tend to have higher T1w/T2w ratio, and ECI signal fraction values compared to males. ASD males, meanwhile, tend to have higher ICI (GM-like) signal fraction values, while in TD participants, age-associated ROIs tend to have higher ICI (GM-like) values in females. Aggregate conduction velocity and aggregate g-ratio do not seem to differ between ASD and TD in their age-by-sex interaction, despite having many significantly different ROIs between ASD and TD participants.

**Figure 9:**
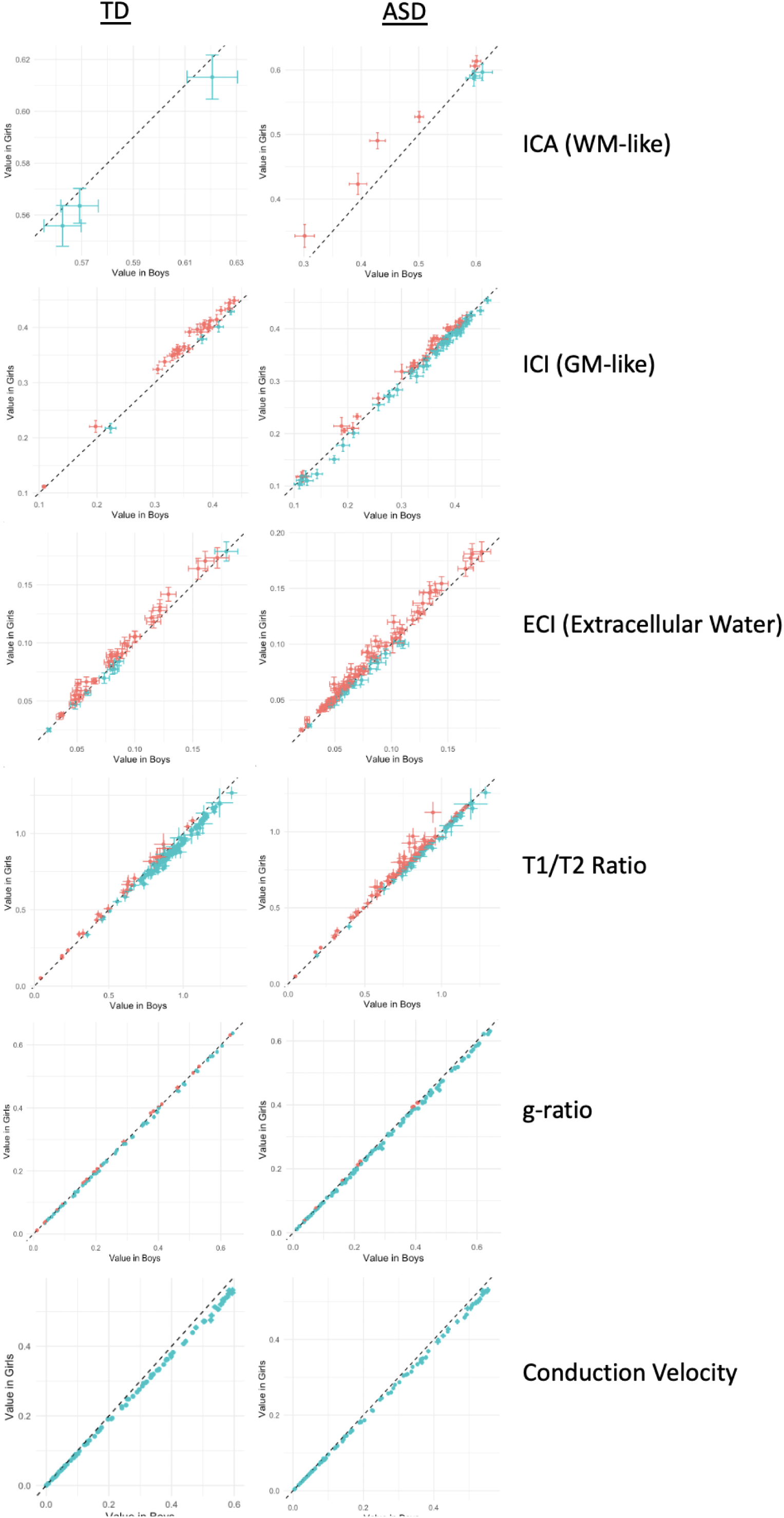
Series of charts displaying mean microstructural values of ROIs significantly associated with age, colored by sex and separated between ASD and TD participants. Diagnosis groups show varying relationships with aging that are not consistent across sexes, with far higher T1W/T2W ratio values in TD boys compared TD girls, while in ASD, girls tend to have higher T1W/T2W ratio and ECI signal fraction values compared to boys. ASD boys meanwhile tend to have higher ICI (GM-like) signal fraction values, while in TD participants age-associated ROIs tend to have higher ICI (GM-like) values in girls. Aggregate conduction velocity and aggregate g-ratio do not seem to differ between ASD and TD in their age*sex interaction.

When microstructural metrics were compared to SCQ total score a similar pattern to diagnosis was observed (Fig. 10). There was a significant positive relationship between ECI signal fraction and SCQ total score in 55 ROIs, indicating that greater extracellular water was associated with a more ASD-like behavioral profile. All but one (the left putamen) were cortical ROIs including the composite cortical ROI. Aggregate g-ratio had a significantly negative association with SCQ total score in 12 ROIs, though this included all 3 ROIs covering the entire corpus callosum. Aggregate conduction velocity however, had a significant negative relationship with SCQ total score in 139 ROIs including both composite cortical and white matter ROIs, indicating that reduced conduction velocity was associated with a more ASD-like behavioral profile (Fig. 11). No other microstructural metric in any ROI was significantly associated with SCQ total score after multiple comparison correction.

**Figure 10:**
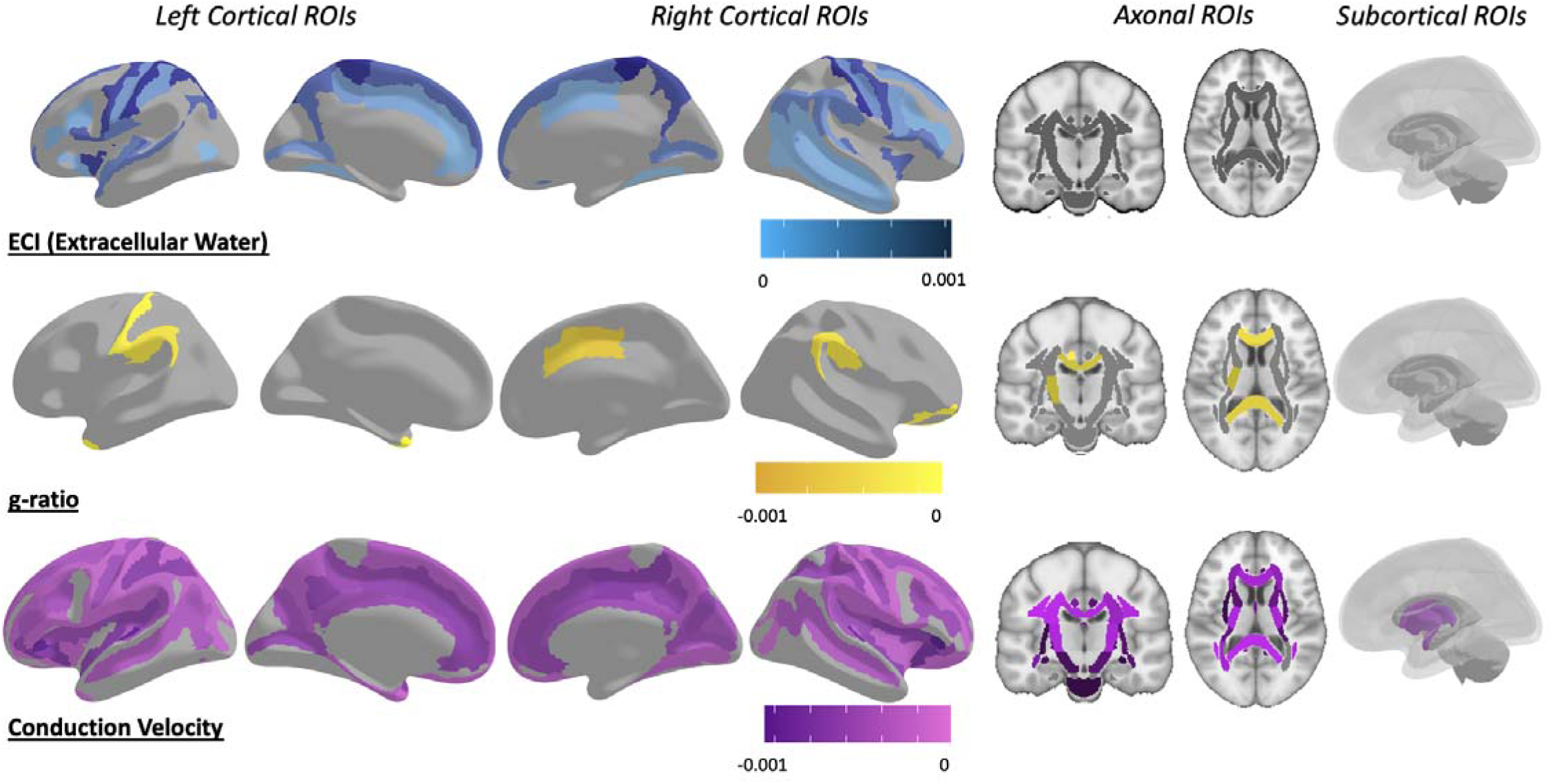
Map of ROIs that were significant at the p<0.05 level between each mean microstructural metric and SCQ total score after multiple comparison corrections. Colored by the slope of the relationship, with extracellular water positively associated, aggregate g-ratio negatively associated, and aggregate conduction velocity negatively associated, in each significant ROI. There was an even more pronounced special specificity in results, with ECI signal fraction ROIs located almost exclusively in cortex, aggregate g-ratio ROIs located across the entire corpus callosum, and aggregate conduction velocity ROIs located throughout the cortex, white matter, and subcortical ROIs.

**Figure 11:**
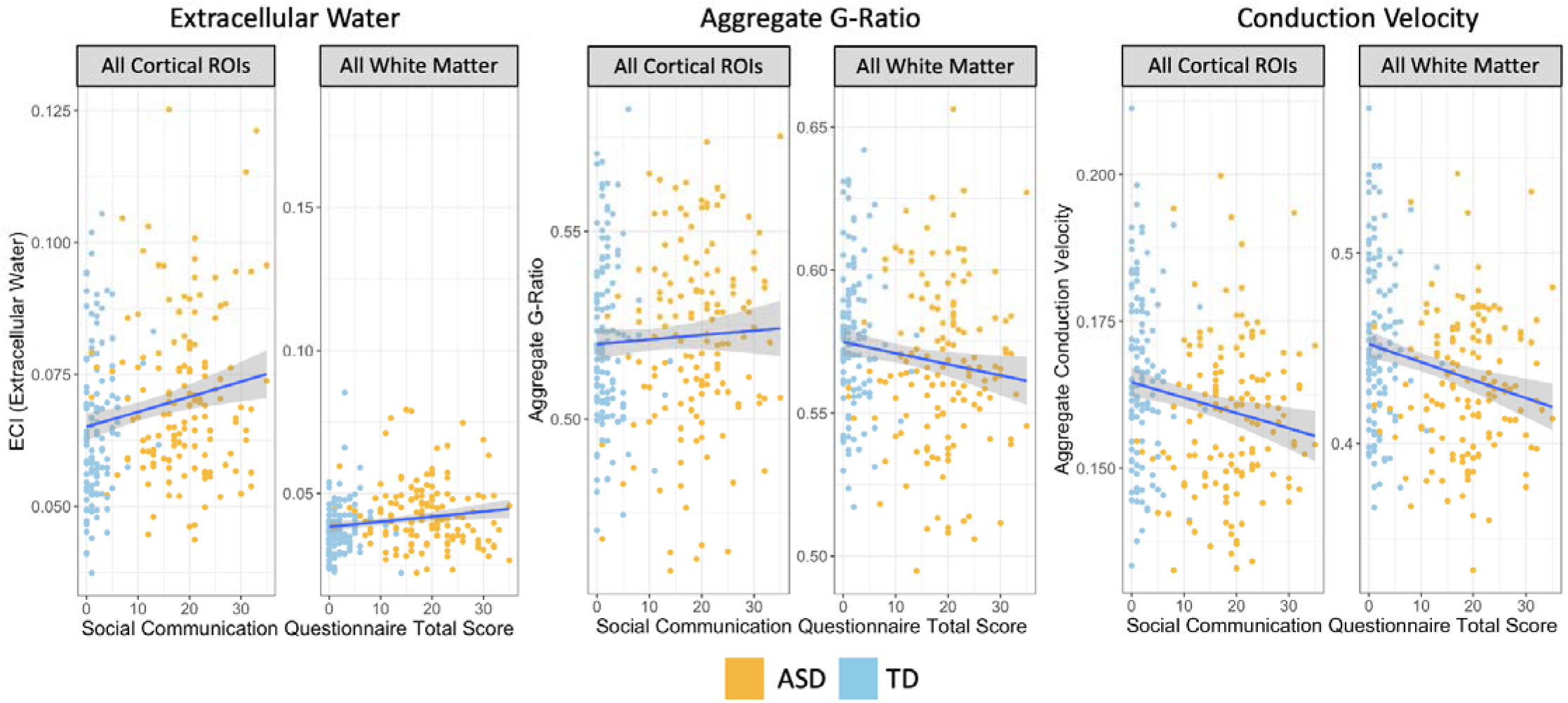
Charts displaying relationship between microstructural metrics with significant ROIs in composite ROIs, colored by ASD diagnosis. The directions of the associations were consistent across all significant ROIs in each microstructural metric. The strongest associations were found in extracellular water metrics in cortical ROIs and aggregate conduction velocity metrics. When examined separately within diagnosis group there was no significant relationship between Social Communication Questionnaire Total Score and brain microstructure in either composite ROI.

## Discussion

This study’s describe subcellular findings that could be the neurobiological basis for reduced long-range connectivity[15–17]. This study reveals widespread significant mean differences between ASD and TD participants while also highlighting significant associations between microstructural metrics and behavioral measures of ASD. These findings were primarily located in the neocortex in the ECI signal fraction, in the axonal WM areas for aggregate g-ratio, and in both neocortex and axonal WM areas for aggregate conduction velocity. ROC analysis showed that aggregate microstructural metrics are highly discriminatory between ASD and TD individuals and are strongly correlated with behavioral metrics.

Multiple studies have previously described consistent increases in cortical thickness[2,4,89] and mean diffusivity with corresponding decreases in FA[90–92] in ASD compared to TD participants. Against this background, the widespread increases of extracellular water in cortical ROIs and decreased aggregate conduction velocity and aggregate g-ratio observed in this study can potentially be interpreted as widespread problems with myelination or as the microstructural result of increased cortical thickness. However, there was no significant difference in T1w/T2w ratio (used as the MVF in this study) between ASD and TD, implying that there is a more complex relationship between myelination and ASD than simply a deficit compared to TD. By weighting intra-axonal volume (measured via the total sum of FDC) into both aggregate g-ratio and aggregate conduction velocity, it is possible that differences are driven by changes to axonal morphology at the sub-cellular level. This may be an architectural difference in cellular microstructure (i.e., axons themselves may be thinner or more dispersed in ASD), or, combined with the increased extracellular water exclusively in the cortex, there is increased active or previous neuroinflammation that stunted axonal and myelin development, leaving a sparser and higher volume cortex having a large portion of that increase simply being extracellular space. This finding is supported by the finding of increased extracellular water and decreased neuronal dispersion by other diffusion models such as NODDI[29]. While the diffusion acquisitions used in this study were not obtained using very high b-values, which could cause extra-axonal signal to be grouped with intra-axonal FDC measurements, there was still significant differences between ASD and TD, suggesting that differences were either large enough to overcome low b-value induced bias or that extra-axonal signal did not meaningfully disrupt measurements. The dedicated extra-axonal signal measurement provided by the ISI signal fraction was not significantly different between ASD and TD in any ROI, suggesting that extra-axonal signal was not meaningfully different between groups. There remain a number of limitations inherent in this work, especially lacking gold-standard *ex-vivo* validation in an animal model. These range from limitations of the CSD framework, for example by enforcing the assumption that all WM fiber diffusion signatures are identical to the WM response function. Deliberately, the g-ratio and conduction velocity values are presented here without alteration or correction to known physiological values. While performing this could have improved accuracy of measurements given the lack of information regarding either metric in an adolescent population it was decided to present all values ‘as-is’, especially since the primary analysis performed is comparative.

Histological work performed on small samples of WM from both TD and ASD adults found that superficial and deep WM had significantly fewer large diameter axonal fibers, more small diameter axonal fibers, and significantly more branching axonal fibers in ASD compared to TD participants[50,93]. Individuals with ASD also had significantly thinner myelin sheaths in all sizes of axon[50]. Though this histological experiment was performed in adults instead of the adolescents in this study, given these two factors, a bias toward smaller diameter axons and thinner myelin, it is likely that the results reported here confirm this prior finding. As illustrated in Fig. 12, the aggregate g-ratio can be lowered if both AVF and MVF are reduced, especially unequally. In absolute terms, it is possible that this adolescent cohort is overall lower than expected in adults due to insufficiently developed myelin and smaller axonal diameters, and thus the TD group can be interpreted as nearer to developed myelin/axonal volumes and thus the initially paradoxical seeming greater g-ratio compared to the ASD group is actually the result of higher axonal diameter[52,81,83,94]. Further comparison of these metrics to age shows that aggregate g-ratio has a negative relationship with age while aggregate conduction velocity has a positive relationship with age during this developmental period (Fig. 8), which may indicate that continued myelin development in both groups (T1w/T2w ratio was not significantly different between groups in any ROI, implying that AVF contributed to significant differences observed here) improves neuronal function during the developmental period with a baseline difference in axonal diameter in the ASD group. The axonal conduction velocity finding has been supported by other studies, and it has been reported that children with ASD have lower conduction velocity using magnetoencephalography[95,96] and in peripheral neurons[97], potentially due to reduced axonal diameter and the prevalence of small fibers. This study reinforces this finding by showing, for the first time noninvasively, widespread reduced aggregate conduction velocity in ASD throughout the brain. The observed reductions in aggregate conduction velocity, in particular, may indicate deficits, particularly in longer-range axonal connections dependent on higher-diameter fibers and myelination. This effect was observed in both aggregate g-ratio and aggregate conduction velocity throughout a wide range of the axonal skeleton, indicating that these functional deficits are systemic. It was further observed that differences between TD and ASD go back to fiber density and cross-section (used as the AVF in the g-ratio and conduction velocity calculations), which, combined with the lack of significance in T1w/T2w ratio measurements, suggests that axonal diameter may be driving the observed differences in the study rather than myelin differences.

**Figure 12:**
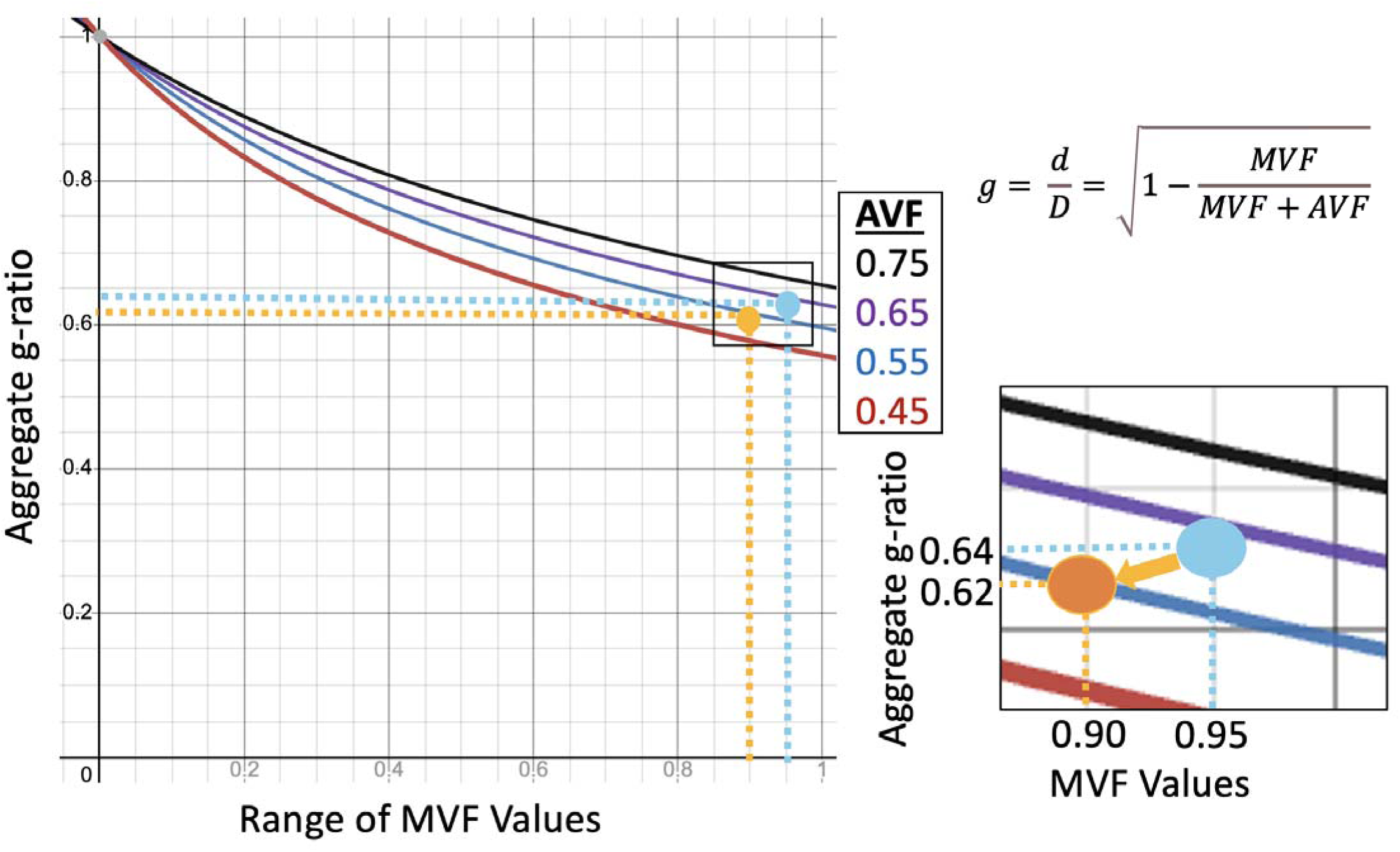
Display relating the aggregate g-ratio equation to ranges of biologically plausible AVF and MVF. In this study we observed a somewhat paradoxical result of lower aggregate g-ratio in ASD (orange circle) in contrast to TD subjects (blue circle), despite significantly lower aggregate conduction velocity and other indicators of microstructural deficits. As illustrated here, for a range of AVF (including a difference of 0.05, observed in the corpus callosum), there is a wide range (nearly 0.1) of reduced MVF values that will nevertheless result in a reduced aggregate g-ratio value. As we observed in this study, this is a plausible explanation to harmonize initially contradictory results of reduced functionality but superior g-ratio.

It should also be noted that extensive work has shown that extracellular water increases in the axonal WM skeleton with age and cerebrovascular injury[68,98–100], but that none of the axonal ROIs tested was significant for ECI signal fraction differences, suggesting that there is no active degeneration of brain cells or microstructure. The findings of extracellular water differences exclusively in cortex pairs with the lack of findings in the axonal signal compartment and in the T1w/T2w ratio measurement of myelin in any ROIs examined. This is also an important reminder that T2-weighted signal and extracellular water are not interchangeable, especially in development. The water in the extracellular space is still close to the bound volume fraction pool water, and the T2-weighted signal was increased enough compared to the T1-weighted signal to serve as a marker of ASD in the T1w/T2w ratio metric. Note, however, that the ECI signal fraction had the largest number of negative correlations with g-ratio and conduction velocity, but, as expected, those metrics were closely related to T1w/T2w ratio in nearly every ROI, despite T1w/T2w ratio subsequently having no ROIs that were different between ASD and TD participants. An additional possibility is that changes in synaptic formation as suggested by several genetic associations between ASD and growth-cone-related genes such as CRMP1 and CRMP2[101]. Recent work using FDC in other model systems, such as the optic nerve, has suggested that fiber cross-section may be a key component of information transfer and downstream neuronal development[102]. Validation studies in animal models or via further histopathological study are required to further investigate the cellular basis of these observed effects.

These alterations in axonal morphology may be responsible for functional changes and subsequent behavioral outcomes. In this study, we observe that reduced aggregate conduction velocity is strongly associated with increased ASD behaviors, as measured by the SCQ. This association is present across a large number of cortical and WM regions, and is very similar to the association between aggregate conduction velocity and ASD diagnosis, suggesting that slowed axonal conduction velocity within and between brain regions could contribute to ASD etiology. Interestingly, this association is present between ASD behavioral metrics and extracellular water in a wide variety of cortical regions but only present between aggregate g-ratio and ASD behavioral metrics in the corpus callosum and a small number of other regions. This matches reports from histological studies where differences in axonal area are much larger than differences in myelination between ASD and TD individuals[93]. This is particularly relevant to our calculation of aggregate conduction velocity which relies on aggregate axonal volume per voxel. Combined with the lack of significant T1w/T2w ratio associations, this suggests that behavioral components of ASD are more likely associated with reduced axonal area and the subsequently slower conduction velocity, than due to a difference in myelination.

## Conclusion

Advanced diffusion and structural microstructure models can characterize a combination of increased extracellular water and decreased aggregate g-ratio and aggregate conduction velocity between ASD and TD participants. Microstructural metrics are similarly associated with a validated measure of ASD behavior, suggesting that particularly slower aggregate conduction velocity relate to higher scores on ASD behavioral evaluations. This study introduces a novel means to calculate aggregate g-ratio using diffusion MRI techniques and a novel calculation of aggregate conduction velocity as well as applying both to the study of ASD for the first time. These findings suggest that differences in axon morphology and subsequently slowed conduction velocity, are widespread throughout the brain and may contribute to the clinical picture of ASD. Altered signal transduction along affected fiber pathways also provides an explanation for recent observations of local versus distal connection dependencies more specifically in those diagnosed with ASD.

## Acknowledgements

This work was performed on behalf of the GENDAAR Consortium (NIH R01 MH100028), and we thank all of our collaborating colleagues, the study participants, and their family members. The funders had no role in study design, data collection and analysis, decision to publish, or preparation of the manuscript. We would like to specifically acknowledge the contributions of Anna Kresse, MPH; Megha Santhosh, MHA; Désirée Lussier-Lévesque, PhD; Emily Neuhaus, PhD; Katy Ankenman, MSW; Jessica Benton, MA; and Rachel Fung, BS.

## Authorship Contributions

BTN conducted the analyses and wrote the manuscript; ZJ curated the data sets used in the analyses; SV contributed to the theoretical concepts; TJD contributed expertise in clinical radiology; JDVH conceived of the overall study, contributed to the theoretical and conceptual considerations, and helped to write the manuscript; KAP contributed expertise in ASD research and provided ACE program leadership (NIH R01 MH100028); SJW, NAK, and JCM provided expertise in ASD clinical assessment and neurophysiological research.

## Supplementary Information

**S1 Table:** Raw means for each neuroimaging metric measured from subjects in each ROI.

**S2 Table:** ECI signal fraction p-values before and after Banjamin and Hochburg multiple comparison adjustment for each of the covariates included in linear model analysis (diagnosis, sex, age, Intracranial volume, and IQ. Significant values at the 0.05 level following multiple comparison adjustment are summarized in Table 1.

**S3 Table:** ICI signal fraction p-values before and after Banjamin and Hochburg multiple comparison adjustment for each of the covariates included in linear model analysis (diagnosis, sex, age, Intracranial volume, and IQ. Significant values at the 0.05 level following multiple comparison adjustment are summarized in Table 1.

**S4 Table:** ICA signal fraction p-values before and after Banjamin and Hochburg multiple comparison adjustment for each of the covariates included in linear model analysis (diagnosis, sex, age, Intracranial volume, and IQ. Significant values at the 0.05 level following multiple comparison adjustment are summarized in Table 1.

**S5 Table:** T1w/T2w ratio p-values before and after Banjamin and Hochburg multiple comparison adjustment for each of the covariates included in linear model analysis (diagnosis, sex, age, Intracranial volume, and IQ. Significant values at the 0.05 level following multiple comparison adjustment are summarized in Table 1.

**S6 Table:** Aggregate g-ratio p-values before and after Banjamin and Hochburg multiple comparison adjustment for each of the covariates included in linear model analysis (diagnosis, sex, age, Intracranial volume, and IQ. Significant values at the 0.05 level following multiple comparison adjustment are summarized in Table 1.

**S7 Table:** Aggregate conduction velocity p-values before and after Banjamin and Hochburg multiple comparison adjustment for each of the covariates included in linear model analysis (diagnosis, sex, age, Intracranial volume, and IQ. Significant values at the 0.05 level following multiple comparison adjustment are summarized in Table 1.

