## Supplementary Table 1 for "Conduction Velocity, G-ratio, and Extracellular Water as Microstructural Characteristics of Autism Spectrum Disorder"

| ROI Name | Region | ICA Signal Fraction | | | | ICI Signal Fraction | | | | ECI Signal Fraction | | | | T1/T2 Ratio | | | | Aggregate G-Ratio | | | | Aggregate Conduction Velocity | | | |
| --- | --- | --- | --- | --- | --- | --- | --- | --- | --- | --- | --- | --- | --- | --- | --- | --- | --- | --- | --- | --- | --- | --- | --- | --- | --- |
|  |  | ASD Mean | ± SD | TD Mean | ± SD | ASD Mean | ± SD | TD Mean | ± SD | ASD Mean | ± SD | TD Mean | ± SD | ASD Mean | ± SD | TD Mean | ± SD | ASD Mean | ± SD | TD Mean | ± SD | ASD Mean | ± SD | TD Mean | ± SD |
| White Matter Composite ROI | Axonal | 0.8343 | 0.0326 | 0.8476 | 0.0270 | 0.1141 | 0.0236 | 0.1099 | 0.0229 | 0.0435 | 0.0161 | 0.0379 | 0.0090 | 1.0459 | 0.1088 | 1.0604 | 0.1108 | 0.5659 | 0.0282 | 0.5748 | 0.0254 | 0.4307 | 0.0352 | 0.4533 | 0.0430 |
| Cortical Composite ROI | Cortical | 0.5109 | 0.0409 | 0.5107 | 0.0366 | 0.3183 | 0.0374 | 0.3212 | 0.0404 | 0.0710 | 0.0158 | 0.0658 | 0.0146 | 0.6908 | 0.1201 | 0.7127 | 0.1125 | 0.5214 | 0.0245 | 0.5208 | 0.0229 | 0.1583 | 0.0128 | 0.1652 | 0.0147 |
| Middle Cerebellar Peduncle | Axonal | 0.8266 | 0.1043 | 0.8301 | 0.1034 | 0.1123 | 0.0924 | 0.1064 | 0.0813 | 0.0579 | 0.0627 | 0.0555 | 0.0788 | 1.1159 | 0.1924 | 1.1038 | 0.1621 | 0.3934 | 0.0318 | 0.4012 | 0.0277 | 0.2853 | 0.0298 | 0.2974 | 0.0312 |
| Pontine Crossing Tract | Axonal | 0.8188 | 0.1162 | 0.8223 | 0.1278 | 0.1097 | 0.0830 | 0.1065 | 0.0800 | 0.0549 | 0.0572 | 0.0570 | 0.0816 | 1.1264 | 0.1620 | 1.1392 | 0.1423 | 0.5553 | 0.0331 | 0.5600 | 0.0284 | 0.4056 | 0.0459 | 0.4232 | 0.0523 |
| Genu of Corpus Callosum | Axonal | 0.8097 | 0.1096 | 0.8250 | 0.0958 | 0.1198 | 0.0903 | 0.1094 | 0.0843 | 0.0618 | 0.0671 | 0.0610 | 0.0729 | 1.1082 | 0.1213 | 1.1183 | 0.1284 | 0.5795 | 0.0412 | 0.5941 | 0.0363 | 0.4538 | 0.0535 | 0.4825 | 0.0591 |
| Body of Corpus Callosum | Axonal | 0.8235 | 0.1099 | 0.8354 | 0.0987 | 0.1115 | 0.0839 | 0.1017 | 0.0784 | 0.0568 | 0.0600 | 0.0576 | 0.0842 | 1.0529 | 0.1120 | 1.0650 | 0.1094 | 0.5994 | 0.0345 | 0.6132 | 0.0314 | 0.4575 | 0.0485 | 0.4877 | 0.0535 |
| Splenium of Corpus Callosum | Axonal | 0.8214 | 0.1037 | 0.8220 | 0.1054 | 0.1140 | 0.0828 | 0.1056 | 0.0778 | 0.0578 | 0.0810 | 0.0680 | 0.1003 | 1.0636 | 0.1190 | 1.0817 | 0.1193 | 0.5713 | 0.0348 | 0.5844 | 0.0335 | 0.4343 | 0.0506 | 0.4658 | 0.0538 |
| Fornix | Axonal | 0.8140 | 0.1157 | 0.8253 | 0.0948 | 0.1159 | 0.0898 | 0.1029 | 0.0762 | 0.0604 | 0.0726 | 0.0629 | 0.0850 | 0.7697 | 0.1180 | 0.7830 | 0.1196 | 0.6395 | 0.0529 | 0.6469 | 0.0643 | 0.3692 | 0.0489 | 0.3851 | 0.0553 |
| Corticospinal Tract R | Axonal | 0.8217 | 0.1086 | 0.8345 | 0.0819 | 0.1158 | 0.0877 | 0.1053 | 0.0723 | 0.0536 | 0.0643 | 0.0570 | 0.0692 | 1.0337 | 0.1818 | 1.0369 | 0.1841 | 0.3916 | 0.0289 | 0.3969 | 0.0231 | 0.3172 | 0.0364 | 0.3258 | 0.0373 |
| Corticospinal Tract L | Axonal | 0.8203 | 0.1057 | 0.8235 | 0.0889 | 0.1147 | 0.0854 | 0.1110 | 0.0798 | 0.0576 | 0.0817 | 0.0629 | 0.0766 | 1.0364 | 0.1801 | 1.0731 | 0.1919 | 0.4283 | 0.0311 | 0.4301 | 0.0302 | 0.3601 | 0.0406 | 0.3730 | 0.0460 |
| Medial Lemniscus R | Axonal | 0.8212 | 0.0974 | 0.8226 | 0.1026 | 0.1165 | 0.0840 | 0.1051 | 0.0778 | 0.0578 | 0.0748 | 0.0638 | 0.0886 | 1.0355 | 0.1234 | 1.0638 | 0.1302 | 0.5808 | 0.0335 | 0.5790 | 0.0345 | 0.4185 | 0.0488 | 0.4295 | 0.0506 |
| Medial Lemniscus L | Axonal | 0.8335 | 0.0963 | 0.8363 | 0.0803 | 0.1051 | 0.0762 | 0.1030 | 0.0729 | 0.0513 | 0.0496 | 0.0563 | 0.0619 | 1.0867 | 0.1209 | 1.1077 | 0.1280 | 0.6220 | 0.0368 | 0.6276 | 0.0388 | 0.4892 | 0.0602 | 0.5080 | 0.0650 |
| Inferior Cerebellar Peduncle R | Axonal | 0.7913 | 0.1622 | 0.8219 | 0.1094 | 0.1178 | 0.0925 | 0.1072 | 0.0776 | 0.0625 | 0.0795 | 0.0664 | 0.1032 | 0.7941 | 0.1726 | 0.8017 | 0.1634 | 0.3851 | 0.0322 | 0.3872 | 0.0295 | 0.2486 | 0.0303 | 0.2554 | 0.0333 |
| Inferior Cerebellar Peduncle L | Axonal | 0.8244 | 0.1214 | 0.8173 | 0.1243 | 0.1103 | 0.0860 | 0.1140 | 0.0880 | 0.0513 | 0.0507 | 0.0642 | 0.1089 | 0.6321 | 0.1040 | 0.6584 | 0.0946 | 0.4222 | 0.0335 | 0.4267 | 0.0292 | 0.2441 | 0.0296 | 0.2534 | 0.0309 |
| Superior Cerebellar Peduncle R | Axonal | 0.8192 | 0.1108 | 0.8284 | 0.1008 | 0.1131 | 0.0858 | 0.1034 | 0.0787 | 0.0569 | 0.0589 | 0.0578 | 0.0770 | 0.8935 | 0.0973 | 0.9103 | 0.0936 | 0.4818 | 0.0308 | 0.4828 | 0.0248 | 0.3339 | 0.0312 | 0.3422 | 0.0349 |
| Superior Cerebellar Peduncle L | Axonal | 0.8176 | 0.1198 | 0.8204 | 0.1172 | 0.1163 | 0.0869 | 0.1052 | 0.0832 | 0.0560 | 0.0668 | 0.0570 | 0.0623 | 0.7769 | 0.0928 | 0.7975 | 0.0912 | 0.4152 | 0.0292 | 0.4179 | 0.0300 | 0.2851 | 0.0280 | 0.2963 | 0.0333 |
| Cerebral Peduncle R | Axonal | 0.8057 | 0.1350 | 0.8321 | 0.0809 | 0.1174 | 0.0922 | 0.1049 | 0.0730 | 0.0647 | 0.0987 | 0.0558 | 0.0572 | 0.8657 | 0.1071 | 0.8816 | 0.1143 | 0.4699 | 0.0257 | 0.4735 | 0.0236 | 0.3142 | 0.0320 | 0.3265 | 0.0351 |
| Cerebral Peduncle L | Axonal | 0.8132 | 0.1245 | 0.8345 | 0.0997 | 0.1156 | 0.0910 | 0.0983 | 0.0736 | 0.0600 | 0.0899 | 0.0580 | 0.0668 | 0.9165 | 0.1126 | 0.9266 | 0.1230 | 0.5208 | 0.0281 | 0.5285 | 0.0291 | 0.3771 | 0.0392 | 0.3935 | 0.0470 |
| Anterior Limb of Internal Capsule R | Axonal | 0.8161 | 0.1190 | 0.8156 | 0.1431 | 0.1145 | 0.0866 | 0.1006 | 0.0773 | 0.0553 | 0.0551 | 0.0645 | 0.0883 | 1.0251 | 0.1070 | 1.0405 | 0.1081 | 0.6284 | 0.0359 | 0.6334 | 0.0328 | 0.4736 | 0.0422 | 0.4890 | 0.0472 |
| Anterior Limb of Internal Capsule L | Axonal | 0.8137 | 0.0953 | 0.8308 | 0.0936 | 0.1174 | 0.0868 | 0.1054 | 0.0784 | 0.0595 | 0.0590 | 0.0605 | 0.0753 | 1.0246 | 0.1125 | 1.0343 | 0.1114 | 0.6381 | 0.0357 | 0.6472 | 0.0312 | 0.4800 | 0.0438 | 0.5015 | 0.0537 |
| Posterior Limb of Internal Capsule R | Axonal | 0.8130 | 0.1198 | 0.8197 | 0.1081 | 0.1145 | 0.0860 | 0.1049 | 0.0823 | 0.0573 | 0.0577 | 0.0633 | 0.0806 | 1.0403 | 0.1233 | 1.0601 | 0.1262 | 0.6581 | 0.0338 | 0.6684 | 0.0299 | 0.5289 | 0.0502 | 0.5598 | 0.0588 |
| Posterior Limb of Internal Capsule L | Axonal | 0.7967 | 0.1530 | 0.8292 | 0.0859 | 0.1220 | 0.0976 | 0.1050 | 0.0797 | 0.0629 | 0.0835 | 0.0628 | 0.0725 | 1.0434 | 0.1164 | 1.0596 | 0.1162 | 0.6652 | 0.0330 | 0.6758 | 0.0319 | 0.5401 | 0.0500 | 0.5717 | 0.0657 |
| Retrolenticular Part of Internal Capsule R | Axonal | 0.8138 | 0.1057 | 0.8274 | 0.0952 | 0.1198 | 0.0899 | 0.1047 | 0.0769 | 0.0586 | 0.0677 | 0.0657 | 0.0878 | 1.1603 | 0.1357 | 1.1819 | 0.1397 | 0.6342 | 0.0353 | 0.6372 | 0.0299 | 0.5337 | 0.0504 | 0.5500 | 0.0553 |
| Retrolenticular Part of Internal Capsule L | Axonal | 0.8168 | 0.1142 | 0.8374 | 0.0826 | 0.1163 | 0.0880 | 0.1035 | 0.0805 | 0.0565 | 0.0666 | 0.0548 | 0.0625 | 1.1667 | 0.1351 | 1.1964 | 0.1356 | 0.6293 | 0.0369 | 0.6304 | 0.0326 | 0.5258 | 0.0488 | 0.5416 | 0.0543 |
| Anterior Corona Radiata R | Axonal | 0.8015 | 0.1122 | 0.8348 | 0.0935 | 0.1216 | 0.0961 | 0.0990 | 0.0795 | 0.0627 | 0.0705 | 0.0589 | 0.0771 | 1.1073 | 0.1186 | 1.1212 | 0.1190 | 0.6000 | 0.0372 | 0.6106 | 0.0333 | 0.4640 | 0.0492 | 0.4871 | 0.0544 |
| Anterior Corona Radiata L | Axonal | 0.8119 | 0.1048 | 0.8193 | 0.0996 | 0.1183 | 0.0929 | 0.1091 | 0.0852 | 0.0588 | 0.0615 | 0.0661 | 0.0837 | 1.1148 | 0.1242 | 1.1334 | 0.1351 | 0.5972 | 0.0374 | 0.6062 | 0.0354 | 0.4569 | 0.0454 | 0.4798 | 0.0544 |
| Superior Corona Radiata R | Axonal | 0.8203 | 0.0962 | 0.8253 | 0.1118 | 0.1177 | 0.0876 | 0.1041 | 0.0853 | 0.0542 | 0.0497 | 0.0646 | 0.0952 | 1.0840 | 0.1204 | 1.1000 | 0.1207 | 0.6571 | 0.0345 | 0.6704 | 0.0309 | 0.5428 | 0.0534 | 0.5778 | 0.0599 |
| Superior Corona Radiata L | Axonal | 0.8291 | 0.0852 | 0.8315 | 0.0886 | 0.1114 | 0.0819 | 0.1059 | 0.0828 | 0.0562 | 0.0551 | 0.0586 | 0.0705 | 1.0860 | 0.1202 | 1.1046 | 0.1230 | 0.6545 | 0.0336 | 0.6649 | 0.0332 | 0.5367 | 0.0505 | 0.5666 | 0.0596 |
| Posterior Corona Radiata R | Axonal | 0.8125 | 0.1126 | 0.8167 | 0.1100 | 0.1178 | 0.0880 | 0.1089 | 0.0836 | 0.0571 | 0.0717 | 0.0697 | 0.1019 | 1.0963 | 0.1203 | 1.1050 | 0.1271 | 0.6412 | 0.0353 | 0.6534 | 0.0316 | 0.5196 | 0.0501 | 0.5467 | 0.0569 |
| Posterior Corona Radiata L | Axonal | 0.8237 | 0.1081 | 0.8248 | 0.1053 | 0.1128 | 0.0819 | 0.1089 | 0.0847 | 0.0585 | 0.0827 | 0.0562 | 0.0665 | 1.0883 | 0.1249 | 1.1066 | 0.1295 | 0.6409 | 0.0356 | 0.6509 | 0.0346 | 0.5136 | 0.0501 | 0.5412 | 0.0537 |
| Posterior Thalamic Radiation R | Axonal | 0.8164 | 0.1120 | 0.8298 | 0.0895 | 0.1150 | 0.0857 | 0.1058 | 0.0811 | 0.0627 | 0.0887 | 0.0601 | 0.0703 | 1.1558 | 0.1366 | 1.1798 | 0.1355 | 0.6357 | 0.0360 | 0.6415 | 0.0319 | 0.5482 | 0.0538 | 0.5737 | 0.0688 |
| Posterior Thalamic Radiation L | Axonal | 0.8088 | 0.1067 | 0.8313 | 0.0964 | 0.1248 | 0.0908 | 0.1039 | 0.0834 | 0.0629 | 0.0790 | 0.0593 | 0.0775 | 1.1945 | 0.1532 | 1.2184 | 0.1589 | 0.6368 | 0.0386 | 0.6375 | 0.0361 | 0.5582 | 0.0553 | 0.5719 | 0.0625 |
| Sagittal Stratum R | Axonal | 0.8282 | 0.1006 | 0.8391 | 0.0799 | 0.1125 | 0.0827 | 0.1047 | 0.0733 | 0.0501 | 0.0482 | 0.0540 | 0.0635 | 1.0948 | 0.1333 | 1.1218 | 0.1259 | 0.5478 | 0.0325 | 0.5520 | 0.0293 | 0.4260 | 0.0395 | 0.4454 | 0.0434 |
| Sagittal Stratum L | Axonal | 0.8032 | 0.1396 | 0.8132 | 0.1396 | 0.1178 | 0.0872 | 0.1070 | 0.0833 | 0.0592 | 0.0624 | 0.0679 | 0.1129 | 1.0324 | 0.1228 | 1.0617 | 0.1229 | 0.5511 | 0.0362 | 0.5540 | 0.0312 | 0.3883 | 0.0395 | 0.4048 | 0.0462 |
| External Capsule R | Axonal | 0.8244 | 0.0973 | 0.8209 | 0.1052 | 0.1100 | 0.0776 | 0.1054 | 0.0810 | 0.0560 | 0.0585 | 0.0698 | 0.0918 | 0.8090 | 0.0873 | 0.8292 | 0.0884 | 0.4775 | 0.0277 | 0.4796 | 0.0237 | 0.2699 | 0.0207 | 0.2798 | 0.0229 |
| External Capsule L | Axonal | 0.8234 | 0.1162 | 0.8235 | 0.1025 | 0.1120 | 0.0811 | 0.1089 | 0.0784 | 0.0581 | 0.0880 | 0.0598 | 0.0758 | 0.8502 | 0.0859 | 0.8691 | 0.0856 | 0.5503 | 0.0293 | 0.5549 | 0.0268 | 0.3336 | 0.0262 | 0.3478 | 0.0308 |
| Cingulum (cingulate gyrus) R | Axonal | 0.8172 | 0.1183 | 0.8225 | 0.1231 | 0.1157 | 0.0844 | 0.1009 | 0.0767 | 0.0532 | 0.0604 | 0.0613 | 0.0720 | 0.8721 | 0.0902 | 0.8850 | 0.0955 | 0.4650 | 0.0356 | 0.4809 | 0.0317 | 0.2825 | 0.0290 | 0.3017 | 0.0313 |
| Cingulum (cingulate gyrus) L | Axonal | 0.8230 | 0.1057 | 0.8326 | 0.0854 | 0.1068 | 0.0811 | 0.1041 | 0.0804 | 0.0559 | 0.0570 | 0.0581 | 0.0654 | 0.8587 | 0.0962 | 0.8760 | 0.1009 | 0.4574 | 0.0321 | 0.4669 | 0.0329 | 0.2785 | 0.0271 | 0.2939 | 0.0322 |
| Cingulum (hippocampus) R | Axonal | 0.8239 | 0.1069 | 0.8279 | 0.0957 | 0.1120 | 0.0834 | 0.1020 | 0.0830 | 0.0520 | 0.0584 | 0.0561 | 0.0621 | 0.9331 | 0.1747 | 0.9229 | 0.1523 | 0.3114 | 0.0298 | 0.3136 | 0.0237 | 0.1708 | 0.0216 | 0.1745 | 0.0242 |
| Cingulum (hippocampus) L | Axonal | 0.8081 | 0.1313 | 0.8224 | 0.1051 | 0.1113 | 0.0834 | 0.1033 | 0.0749 | 0.0680 | 0.0834 | 0.0677 | 0.0969 | 0.9697 | 0.1875 | 0.9845 | 0.2274 | 0.2572 | 0.0269 | 0.2609 | 0.0259 | 0.1334 | 0.0164 | 0.1362 | 0.0172 |
| Fornix / Stria terminalis R | Axonal | 0.8166 | 0.1258 | 0.8221 | 0.0916 | 0.1100 | 0.0858 | 0.1104 | 0.0808 | 0.0577 | 0.0691 | 0.0622 | 0.0788 | 0.8042 | 0.0925 | 0.8219 | 0.0935 | 0.5370 | 0.0367 | 0.5457 | 0.0286 | 0.2903 | 0.0278 | 0.3029 | 0.0264 |
| Fornix / Stria terminalis L | Axonal | 0.8109 | 0.1240 | 0.8360 | 0.0892 | 0.1130 | 0.0887 | 0.1021 | 0.0809 | 0.0581 | 0.0714 | 0.0583 | 0.0713 | 0.7820 | 0.0848 | 0.8012 | 0.0883 | 0.5465 | 0.0387 | 0.5517 | 0.0326 | 0.2951 | 0.0293 | 0.3055 | 0.0289 |
| Superior Longitudinal Fasciculus R | Axonal | 0.8044 | 0.1537 | 0.8221 | 0.0960 | 0.1126 | 0.0837 | 0.1107 | 0.0896 | 0.0562 | 0.0665 | 0.0611 | 0.0739 | 1.0913 | 0.1204 | 1.1151 | 0.1268 | 0.6108 | 0.0342 | 0.6221 | 0.0313 | 0.4945 | 0.0511 | 0.5274 | 0.0593 |
| Superior Longitudinal Fasciculus L | Axonal | 0.8297 | 0.0974 | 0.8240 | 0.0983 | 0.1110 | 0.0833 | 0.1144 | 0.0898 | 0.0549 | 0.0596 | 0.0598 | 0.0778 | 1.1155 | 0.1258 | 1.1349 | 0.1312 | 0.6211 | 0.0347 | 0.6317 | 0.0353 | 0.5093 | 0.0521 | 0.5389 | 0.0633 |
| Superior Fronto-Occipital Fasciculus R | Axonal | 0.8253 | 0.1017 | 0.8187 | 0.1139 | 0.1125 | 0.0773 | 0.1044 | 0.0803 | 0.0565 | 0.0608 | 0.0665 | 0.0963 | 1.0927 | 0.1180 | 1.1027 | 0.1167 | 0.6503 | 0.0369 | 0.6594 | 0.0320 | 0.5326 | 0.0542 | 0.5543 | 0.0561 |
| Superior Fronto-Occipital Fasciculus L | Axonal | 0.8119 | 0.1152 | 0.8143 | 0.1337 | 0.1154 | 0.0811 | 0.1034 | 0.0748 | 0.0591 | 0.0655 | 0.0610 | 0.0735 | 1.1094 | 0.1195 | 1.1210 | 0.1285 | 0.6463 | 0.0351 | 0.6513 | 0.0338 | 0.5336 | 0.0508 | 0.5490 | 0.0609 |
| Uncinate Fasciculus R | Axonal | 0.8154 | 0.1033 | 0.8238 | 0.1035 | 0.1197 | 0.0912 | 0.1091 | 0.0819 | 0.0570 | 0.0599 | 0.0595 | 0.0829 | 0.8420 | 0.1011 | 0.8605 | 0.1082 | 0.4610 | 0.0397 | 0.4590 | 0.0422 | 0.2848 | 0.0357 | 0.2907 | 0.0406 |
| Uncinate Fasciculus L | Axonal | 0.8234 | 0.1017 | 0.8258 | 0.1098 | 0.1125 | 0.0842 | 0.1050 | 0.0846 | 0.0530 | 0.0574 | 0.0576 | 0.0701 | 0.9066 | 0.1145 | 0.9262 | 0.1114 | 0.5627 | 0.0442 | 0.5689 | 0.0364 | 0.3701 | 0.0477 | 0.3881 | 0.0534 |
| Tapetum R | Axonal | 0.8108 | 0.1362 | 0.8235 | 0.1171 | 0.1103 | 0.0821 | 0.1029 | 0.0784 | 0.0545 | 0.0575 | 0.0636 | 0.0835 | 1.1084 | 0.1347 | 1.1313 | 0.1286 | 0.6307 | 0.0372 | 0.6378 | 0.0327 | 0.5014 | 0.0473 | 0.5257 | 0.0537 |
| Tapetum L | Axonal | 0.8161 | 0.1063 | 0.8242 | 0.0930 | 0.1176 | 0.0886 | 0.1070 | 0.0851 | 0.0595 | 0.0654 | 0.0647 | 0.0758 | 0.7254 | 0.0994 | 0.7485 | 0.0973 | 0.5293 | 0.0662 | 0.5233 | 0.0618 | 0.2923 | 0.0371 | 0.3050 | 0.0396 |
| Left-Accumbens-area | Sub-Cortical | 0.6248 | 0.0741 | 0.6367 | 0.0711 | 0.3433 | 0.0699 | 0.3373 | 0.0679 | 0.0317 | 0.0133 | 0.0261 | 0.0113 | 0.9174 | 0.1211 | 0.9349 | 0.1085 | 0.5901 | 0.0543 | 0.5993 | 0.0515 | 0.3972 | 0.0717 | 0.4162 | 0.0755 |
| Left-Amygdala | Sub-Cortical | 0.6134 | 0.0672 | 0.6062 | 0.0605 | 0.2828 | 0.0553 | 0.2997 | 0.0613 | 0.1026 | 0.0426 | 0.0940 | 0.0260 | 0.9404 | 0.1293 | 0.9536 | 0.1102 | 0.5810 | 0.0367 | 0.5829 | 0.0307 | 0.3911 | 0.0399 | 0.4029 | 0.0461 |
| Left-Caudate | Sub-Cortical | 0.6910 | 0.0701 | 0.7132 | 0.0586 | 0.1883 | 0.0514 | 0.1846 | 0.0435 | 0.1200 | 0.0827 | 0.1017 | 0.0678 | 0.8689 | 0.0969 | 0.8901 | 0.0961 | 0.5759 | 0.0330 | 0.5815 | 0.0299 | 0.3430 | 0.0280 | 0.3569 | 0.0305 |
| Left-Cerebellum-Cortex | Other | 0.4577 | 0.1165 | 0.4607 | 0.0979 | 0.1878 | 0.0506 | 0.1926 | 0.0461 | 0.0397 | 0.0206 | 0.0366 | 0.0170 | 0.4528 | 0.0738 | 0.4584 | 0.0639 | 0.2197 | 0.0141 | 0.2200 | 0.0124 | 0.1435 | 0.0133 | 0.1471 | 0.0139 |
| Left-Hippocampus | Sub-Cortical | 0.5606 | 0.0565 | 0.5549 | 0.0539 | 0.3246 | 0.0541 | 0.3434 | 0.0581 | 0.1144 | 0.0426 | 0.1016 | 0.0290 | 0.8660 | 0.1014 | 0.8723 | 0.0914 | 0.4774 | 0.0324 | 0.4783 | 0.0288 | 0.2759 | 0.0297 | 0.2821 | 0.0321 |
| Left-Putamen | Sub-Cortical | 0.7592 | 0.0568 | 0.7736 | 0.0467 | 0.2246 | 0.0541 | 0.2123 | 0.0452 | 0.0163 | 0.0047 | 0.0141 | 0.0034 | 0.9032 | 0.0901 | 0.9208 | 0.0907 | 0.5956 | 0.0317 | 0.6005 | 0.0273 | 0.4010 | 0.0335 | 0.4174 | 0.0366 |
| Left-Thalamus-Proper | Sub-Cortical | 0.7180 | 0.0579 | 0.7319 | 0.0467 | 0.2024 | 0.0427 | 0.1994 | 0.0380 | 0.0795 | 0.0459 | 0.0686 | 0.0296 | 0.8990 | 0.0961 | 0.9233 | 0.1038 | 0.5772 | 0.0301 | 0.5834 | 0.0281 | 0.3576 | 0.0279 | 0.3751 | 0.0365 |
| Right-Accumbens-area | Sub-Cortical | 0.6496 | 0.0742 | 0.6596 | 0.0671 | 0.3195 | 0.0687 | 0.3131 | 0.0642 | 0.0307 | 0.0127 | 0.0270 | 0.0140 | 0.8688 | 0.1022 | 0.8705 | 0.1002 | 0.5458 | 0.0521 | 0.5614 | 0.0502 | 0.3433 | 0.0570 | 0.3607 | 0.0641 |
| Right-Amygdala | Sub-Cortical | 0.5898 | 0.0676 | 0.5837 | 0.0621 | 0.3049 | 0.0574 | 0.3182 | 0.0602 | 0.1051 | 0.0393 | 0.0956 | 0.0289 | 0.9250 | 0.1223 | 0.9610 | 0.1312 | 0.5452 | 0.0314 | 0.5441 | 0.0300 | 0.3564 | 0.0331 | 0.3656 | 0.0414 |
| Right-Caudate | Sub-Cortical | 0.6957 | 0.0669 | 0.7152 | 0.0509 | 0.2155 | 0.0531 | 0.2130 | 0.0458 | 0.0885 | 0.0711 | 0.0716 | 0.0476 | 0.8707 | 0.0976 | 0.8967 | 0.0948 | 0.5702 | 0.0344 | 0.5701 | 0.0308 | 0.3312 | 0.0275 | 0.3412 | 0.0263 |
| Right-Cerebellum-Cortex | Other | 0.4544 | 0.1212 | 0.4628 | 0.1080 | 0.1804 | 0.0512 | 0.1819 | 0.0470 | 0.0361 | 0.0186 | 0.0337 | 0.0173 | 0.4950 | 0.0792 | 0.4996 | 0.0732 | 0.2313 | 0.0157 | 0.2334 | 0.0154 | 0.1481 | 0.0142 | 0.1540 | 0.0171 |
| Right-Hippocampus | Sub-Cortical | 0.5496 | 0.0564 | 0.5421 | 0.0540 | 0.3319 | 0.0550 | 0.3460 | 0.0569 | 0.1182 | 0.0450 | 0.1110 | 0.0326 | 0.8749 | 0.1028 | 0.8997 | 0.1052 | 0.4570 | 0.0283 | 0.4577 | 0.0264 | 0.2647 | 0.0269 | 0.2742 | 0.0311 |
| Right-Putamen | Sub-Cortical | 0.7376 | 0.0626 | 0.7500 | 0.0547 | 0.2430 | 0.0588 | 0.2325 | 0.0514 | 0.0194 | 0.0058 | 0.0175 | 0.0051 | 0.9139 | 0.1015 | 0.9381 | 0.1042 | 0.5626 | 0.0341 | 0.5648 | 0.0312 | 0.3764 | 0.0349 | 0.3910 | 0.0393 |
| Right-Thalamus-Proper | Sub-Cortical | 0.7298 | 0.0594 | 0.7411 | 0.0514 | 0.1993 | 0.0447 | 0.1960 | 0.0429 | 0.0709 | 0.0448 | 0.0630 | 0.0319 | 0.8896 | 0.0861 | 0.9056 | 0.0898 | 0.5800 | 0.0293 | 0.5868 | 0.0259 | 0.3553 | 0.0267 | 0.3700 | 0.0333 |
| ctx_lh_G_Ins_lg_and_S_cent_ins | Cortical | 0.4466 | 0.0514 | 0.4458 | 0.0514 | 0.3957 | 0.0520 | 0.4070 | 0.0562 | 0.1565 | 0.0468 | 0.1463 | 0.0395 | 0.5641 | 0.0834 | 0.5701 | 0.0758 | 0.1729 | 0.0232 | 0.1740 | 0.0208 | 0.0687 | 0.0089 | 0.0702 | 0.0091 |
| ctx_lh_G_and_S_cingul-Ant | Cortical | 0.6035 | 0.0540 | 0.6053 | 0.0485 | 0.3512 | 0.0522 | 0.3541 | 0.0460 | 0.0451 | 0.0134 | 0.0404 | 0.0137 | 0.9035 | 0.1088 | 0.9252 | 0.1156 | 0.4288 | 0.0296 | 0.4360 | 0.0296 | 0.3043 | 0.0345 | 0.3183 | 0.0397 |
| ctx_lh_G_and_S_cingul-Mid-Ant | Cortical | 0.5994 | 0.0557 | 0.6029 | 0.0558 | 0.3660 | 0.0514 | 0.3626 | 0.0510 | 0.0346 | 0.0111 | 0.0294 | 0.0115 | 0.9141 | 0.1065 | 0.9299 | 0.1089 | 0.5121 | 0.0313 | 0.5200 | 0.0305 | 0.3824 | 0.0381 | 0.3999 | 0.0455 |
| ctx_lh_G_and_S_cingul-Mid-Post | Cortical | 0.5716 | 0.0627 | 0.5899 | 0.0605 | 0.3931 | 0.0573 | 0.3772 | 0.0581 | 0.0352 | 0.0138 | 0.0293 | 0.0130 | 0.8947 | 0.1018 | 0.9141 | 0.1030 | 0.4872 | 0.0303 | 0.4970 | 0.0276 | 0.3534 | 0.0354 | 0.3713 | 0.0389 |
| ctx_lh_G_and_S_frontomargin | Cortical | 0.3649 | 0.1227 | 0.3513 | 0.1240 | 0.1634 | 0.0706 | 0.1668 | 0.0681 | 0.0636 | 0.0312 | 0.0540 | 0.0254 | 0.8874 | 0.3944 | 0.8866 | 0.3975 | 0.1949 | 0.0236 | 0.1942 | 0.0233 | 0.0876 | 0.0121 | 0.0914 | 0.0153 |
| ctx_lh_G_and_S_occipital_inf | Cortical | 0.5161 | 0.0953 | 0.5051 | 0.0867 | 0.3169 | 0.1025 | 0.3180 | 0.0972 | 0.0927 | 0.0449 | 0.0885 | 0.0372 | 0.3129 | 0.2150 | 0.3360 | 0.2006 | 0.0202 | 0.0069 | 0.0215 | 0.0067 | 0.0066 | 0.0024 | 0.0073 | 0.0028 |
| ctx_lh_G_and_S_paracentral | Cortical | 0.3315 | 0.1081 | 0.3220 | 0.1127 | 0.1967 | 0.0536 | 0.1852 | 0.0578 | 0.1320 | 0.0806 | 0.1233 | 0.0831 | 0.3663 | 0.0983 | 0.3706 | 0.0764 | 0.1769 | 0.0117 | 0.1773 | 0.0113 | 0.1170 | 0.0139 | 0.1212 | 0.0144 |
| ctx_lh_G_and_S_subcentral | Cortical | 0.4885 | 0.0668 | 0.4846 | 0.0566 | 0.4156 | 0.0596 | 0.4295 | 0.0554 | 0.0941 | 0.0408 | 0.0852 | 0.0304 | 0.6876 | 0.1652 | 0.7144 | 0.1629 | 0.1506 | 0.0145 | 0.1528 | 0.0133 | 0.0819 | 0.0091 | 0.0862 | 0.0104 |
| ctx_lh_G_and_S_transv_frontopol | Cortical | 0.4130 | 0.1900 | 0.3846 | 0.1998 | 0.1332 | 0.0825 | 0.1310 | 0.0864 | 0.0858 | 0.0568 | 0.0743 | 0.0508 | 0.8538 | 0.4728 | 0.8979 | 0.5118 | 0.1308 | 0.0241 | 0.1326 | 0.0259 | 0.0544 | 0.0105 | 0.0581 | 0.0107 |
| ctx_lh_G_cingul-Post-dorsal | Cortical | 0.5561 | 0.0731 | 0.5696 | 0.0731 | 0.4235 | 0.0695 | 0.4133 | 0.0708 | 0.0204 | 0.0097 | 0.0171 | 0.0099 | 0.7725 | 0.0819 | 0.7819 | 0.0914 | 0.4179 | 0.0363 | 0.4265 | 0.0314 | 0.2024 | 0.0230 | 0.2115 | 0.0221 |
| ctx_lh_G_cingul-Post-ventral | Cortical | 0.5760 | 0.0994 | 0.5743 | 0.1147 | 0.3227 | 0.0780 | 0.3274 | 0.0858 | 0.1012 | 0.0816 | 0.0983 | 0.1017 | 0.6566 | 0.0893 | 0.6591 | 0.0850 | 0.1676 | 0.0293 | 0.1754 | 0.0266 | 0.0801 | 0.0156 | 0.0832 | 0.0158 |
| ctx_lh_G_cuneus | Cortical | 0.4849 | 0.0515 | 0.4852 | 0.0548 | 0.3933 | 0.0488 | 0.3972 | 0.0477 | 0.1044 | 0.0340 | 0.1024 | 0.0378 | 0.8907 | 0.1531 | 0.9279 | 0.1592 | 0.2748 | 0.0218 | 0.2777 | 0.0259 | 0.1636 | 0.0226 | 0.1710 | 0.0274 |
| ctx_lh_G_front_inf-Opercular | Cortical | 0.4851 | 0.0483 | 0.4821 | 0.0428 | 0.4152 | 0.0504 | 0.4309 | 0.0503 | 0.0986 | 0.0424 | 0.0863 | 0.0318 | 0.8282 | 0.2159 | 0.8975 | 0.2349 | 0.2400 | 0.0198 | 0.2420 | 0.0180 | 0.1211 | 0.0133 | 0.1257 | 0.0140 |
| ctx_lh_G_front_inf-Orbital | Cortical | 0.5643 | 0.0788 | 0.5545 | 0.0731 | 0.3389 | 0.0660 | 0.3444 | 0.0684 | 0.0655 | 0.0327 | 0.0603 | 0.0334 | 0.7561 | 0.1507 | 0.7888 | 0.1508 | 0.3577 | 0.0324 | 0.3621 | 0.0299 | 0.2107 | 0.0286 | 0.2258 | 0.0374 |
| ctx_lh_G_front_inf-Triangul | Cortical | 0.5033 | 0.0538 | 0.5060 | 0.0427 | 0.3966 | 0.0580 | 0.4016 | 0.0491 | 0.0895 | 0.0362 | 0.0831 | 0.0321 | 0.7541 | 0.2483 | 0.8421 | 0.2667 | 0.2195 | 0.0252 | 0.2173 | 0.0217 | 0.1148 | 0.0166 | 0.1194 | 0.0156 |
| ctx_lh_G_front_middle | Cortical | 0.4855 | 0.0475 | 0.4837 | 0.0494 | 0.4090 | 0.0517 | 0.4098 | 0.0541 | 0.0968 | 0.0416 | 0.0898 | 0.0315 | 0.7360 | 0.3087 | 0.8305 | 0.3030 | 0.1696 | 0.0216 | 0.1686 | 0.0194 | 0.0784 | 0.0113 | 0.0833 | 0.0105 |
| ctx_lh_G_front_sup | Cortical | 0.4979 | 0.0533 | 0.4856 | 0.0729 | 0.3302 | 0.0464 | 0.3245 | 0.0622 | 0.1113 | 0.0370 | 0.0996 | 0.0369 | 0.5841 | 0.1623 | 0.6238 | 0.1525 | 0.2645 | 0.0164 | 0.2669 | 0.0156 | 0.1642 | 0.0160 | 0.1717 | 0.0177 |
| ctx_lh_G_insular_short | Cortical | 0.4550 | 0.0481 | 0.4665 | 0.0468 | 0.3973 | 0.0492 | 0.4078 | 0.0448 | 0.1466 | 0.0611 | 0.1247 | 0.0466 | 0.5839 | 0.0992 | 0.6091 | 0.0951 | 0.1987 | 0.0281 | 0.1983 | 0.0302 | 0.0821 | 0.0117 | 0.0848 | 0.0139 |
| ctx_lh_G_oc-temp_lat-fusifor | Cortical | 0.5223 | 0.0675 | 0.5112 | 0.0599 | 0.3915 | 0.0606 | 0.4011 | 0.0553 | 0.0776 | 0.0256 | 0.0815 | 0.0251 | 0.8071 | 0.1396 | 0.8146 | 0.1063 | 0.4473 | 0.0335 | 0.4429 | 0.0321 | 0.2235 | 0.0227 | 0.2234 | 0.0224 |
| ctx_lh_G_oc-temp_med-Lingual | Cortical | 0.5052 | 0.0580 | 0.5043 | 0.0591 | 0.4014 | 0.0573 | 0.4065 | 0.0586 | 0.0862 | 0.0277 | 0.0843 | 0.0330 | 0.8208 | 0.1407 | 0.8396 | 0.1285 | 0.2096 | 0.0185 | 0.2131 | 0.0191 | 0.1023 | 0.0114 | 0.1057 | 0.0103 |
| ctx_lh_G_oc-temp_med-Parahip | Cortical | 0.5691 | 0.0800 | 0.5548 | 0.0688 | 0.3171 | 0.0718 | 0.3458 | 0.0678 | 0.0854 | 0.0323 | 0.0814 | 0.0326 | 1.1789 | 0.2549 | 1.1829 | 0.2377 | 0.2210 | 0.0229 | 0.2247 | 0.0201 | 0.1462 | 0.0171 | 0.1517 | 0.0177 |
| ctx_lh_G_occipital_middle | Cortical | 0.4817 | 0.1014 | 0.4802 | 0.0820 | 0.3089 | 0.1156 | 0.3236 | 0.0989 | 0.0940 | 0.0388 | 0.0878 | 0.0343 | 0.6063 | 0.3662 | 0.6454 | 0.3363 | 0.0785 | 0.0172 | 0.0802 | 0.0179 | 0.0318 | 0.0085 | 0.0355 | 0.0089 |
| ctx_lh_G_occipital_sup | Cortical | 0.4356 | 0.1459 | 0.4141 | 0.1358 | 0.1873 | 0.1172 | 0.1952 | 0.1077 | 0.1274 | 0.0664 | 0.1237 | 0.0647 | 0.7635 | 0.4097 | 0.8352 | 0.4200 | 0.1267 | 0.0251 | 0.1273 | 0.0242 | 0.0531 | 0.0111 | 0.0568 | 0.0116 |
| ctx_lh_G_orbital | Cortical | 0.3868 | 0.1055 | 0.3630 | 0.1072 | 0.1778 | 0.0795 | 0.1755 | 0.0766 | 0.0779 | 0.0241 | 0.0720 | 0.0236 | 0.6795 | 0.1576 | 0.7044 | 0.1567 | 0.0774 | 0.0115 | 0.0778 | 0.0097 | 0.0404 | 0.0061 | 0.0422 | 0.0062 |
| ctx_lh_G_pariet_inf-Angular | Cortical | 0.5067 | 0.0808 | 0.4936 | 0.0746 | 0.3349 | 0.0948 | 0.3589 | 0.0937 | 0.1264 | 0.0417 | 0.1172 | 0.0445 | 0.6238 | 0.3068 | 0.6580 | 0.3033 | 0.1371 | 0.0214 | 0.1380 | 0.0204 | 0.0599 | 0.0117 | 0.0636 | 0.0109 |
| ctx_lh_G_pariet_inf-Supramar | Cortical | 0.4715 | 0.0553 | 0.4764 | 0.0514 | 0.4412 | 0.0588 | 0.4429 | 0.0545 | 0.0846 | 0.0337 | 0.0788 | 0.0326 | 0.6611 | 0.2423 | 0.6893 | 0.2326 | 0.1699 | 0.0179 | 0.1741 | 0.0187 | 0.0788 | 0.0118 | 0.0836 | 0.0115 |
| ctx_lh_G_parietal_sup | Cortical | 0.4197 | 0.0944 | 0.3997 | 0.0898 | 0.2734 | 0.0840 | 0.2825 | 0.0910 | 0.1760 | 0.0706 | 0.1723 | 0.0747 | 0.4308 | 0.1691 | 0.4494 | 0.1644 | 0.1211 | 0.0150 | 0.1231 | 0.0164 | 0.0535 | 0.0081 | 0.0560 | 0.0078 |
| ctx_lh_G_postcentral | Cortical | 0.3857 | 0.0654 | 0.3761 | 0.0712 | 0.3267 | 0.0698 | 0.3274 | 0.0693 | 0.1666 | 0.0588 | 0.1594 | 0.0687 | 0.3027 | 0.1485 | 0.3202 | 0.1390 | 0.0449 | 0.0085 | 0.0471 | 0.0093 | 0.0205 | 0.0038 | 0.0227 | 0.0042 |
| ctx_lh_G_precentral | Cortical | 0.4111 | 0.0645 | 0.3954 | 0.0684 | 0.3429 | 0.0576 | 0.3392 | 0.0664 | 0.1436 | 0.0463 | 0.1330 | 0.0526 | 0.3877 | 0.1531 | 0.4257 | 0.1498 | 0.0715 | 0.0087 | 0.0735 | 0.0096 | 0.0375 | 0.0054 | 0.0407 | 0.0053 |
| ctx_lh_G_precuneus | Cortical | 0.4752 | 0.0470 | 0.4803 | 0.0493 | 0.4077 | 0.0494 | 0.4084 | 0.0514 | 0.1049 | 0.0399 | 0.0954 | 0.0358 | 0.7906 | 0.1107 | 0.7976 | 0.0969 | 0.3487 | 0.0239 | 0.3547 | 0.0243 | 0.1878 | 0.0196 | 0.1978 | 0.0222 |
| ctx_lh_G_rectus | Cortical | 0.4258 | 0.1343 | 0.4182 | 0.1262 | 0.1639 | 0.0795 | 0.1679 | 0.0838 | 0.0726 | 0.0344 | 0.0722 | 0.0297 | 1.0782 | 0.3243 | 1.1360 | 0.3920 | 0.1110 | 0.0137 | 0.1131 | 0.0139 | 0.0642 | 0.0080 | 0.0673 | 0.0100 |
| ctx_lh_G_subcallosal | Cortical | 0.5236 | 0.0779 | 0.5269 | 0.0794 | 0.4029 | 0.0750 | 0.4064 | 0.0724 | 0.0683 | 0.0349 | 0.0622 | 0.0332 | 0.9858 | 0.2117 | 0.9788 | 0.1810 | 0.2775 | 0.0560 | 0.2910 | 0.0569 | 0.1683 | 0.0537 | 0.1797 | 0.0536 |
| ctx_lh_G_temp_sup-G_T_transv | Cortical | 0.4800 | 0.0750 | 0.4945 | 0.0858 | 0.4093 | 0.0606 | 0.4105 | 0.0719 | 0.1104 | 0.0499 | 0.0947 | 0.0469 | 0.8602 | 0.0937 | 0.8778 | 0.0963 | 0.4062 | 0.0254 | 0.4116 | 0.0288 | 0.2375 | 0.0256 | 0.2495 | 0.0298 |
| ctx_lh_G_temp_sup-Lateral | Cortical | 0.4758 | 0.0529 | 0.4735 | 0.0476 | 0.4008 | 0.0545 | 0.4074 | 0.0577 | 0.1047 | 0.0403 | 0.0987 | 0.0363 | 0.6040 | 0.2394 | 0.6646 | 0.2435 | 0.0952 | 0.0118 | 0.0934 | 0.0108 | 0.0468 | 0.0066 | 0.0484 | 0.0061 |
| ctx_lh_G_temp_sup-Plan_polar | Cortical | 0.5274 | 0.0882 | 0.5274 | 0.0721 | 0.3253 | 0.0612 | 0.3359 | 0.0554 | 0.1416 | 0.0659 | 0.1354 | 0.0624 | 0.8646 | 0.1203 | 0.8995 | 0.1158 | 0.5717 | 0.0406 | 0.5723 | 0.0414 | 0.3753 | 0.0459 | 0.3944 | 0.0560 |
| ctx_lh_G_temp_sup-Plan_tempo | Cortical | 0.4878 | 0.0617 | 0.4917 | 0.0608 | 0.4372 | 0.0583 | 0.4366 | 0.0595 | 0.0747 | 0.0344 | 0.0715 | 0.0350 | 0.8385 | 0.2679 | 0.8721 | 0.2536 | 0.3454 | 0.0276 | 0.3493 | 0.0283 | 0.2077 | 0.0262 | 0.2194 | 0.0302 |
| ctx_lh_G_temporal_inf | Cortical | 0.5536 | 0.0860 | 0.5426 | 0.0735 | 0.2803 | 0.0651 | 0.2966 | 0.0640 | 0.0587 | 0.0188 | 0.0589 | 0.0171 | 0.5237 | 0.2331 | 0.5550 | 0.2091 | 0.0432 | 0.0082 | 0.0438 | 0.0082 | 0.0217 | 0.0045 | 0.0222 | 0.0042 |
| ctx_lh_G_temporal_middle | Cortical | 0.5072 | 0.0572 | 0.4980 | 0.0595 | 0.4187 | 0.0560 | 0.4291 | 0.0596 | 0.0605 | 0.0195 | 0.0582 | 0.0185 | 0.4639 | 0.2162 | 0.5012 | 0.2135 | 0.0749 | 0.0127 | 0.0746 | 0.0109 | 0.0332 | 0.0062 | 0.0350 | 0.0062 |
| ctx_lh_Lat_Fis-ant-Horizont | Cortical | 0.7124 | 0.0847 | 0.7421 | 0.0703 | 0.2545 | 0.0752 | 0.2318 | 0.0635 | 0.0329 | 0.0163 | 0.0259 | 0.0122 | 0.9815 | 0.1192 | 1.0009 | 0.1192 | 0.5723 | 0.0393 | 0.5815 | 0.0406 | 0.4234 | 0.0541 | 0.4480 | 0.0645 |
| ctx_lh_Lat_Fis-ant-Vertical | Cortical | 0.6381 | 0.1246 | 0.6324 | 0.1211 | 0.3195 | 0.1077 | 0.3283 | 0.1077 | 0.0424 | 0.0289 | 0.0393 | 0.0265 | 0.9387 | 0.1154 | 0.9455 | 0.1185 | 0.5760 | 0.0439 | 0.5847 | 0.0401 | 0.4027 | 0.0575 | 0.4131 | 0.0594 |
| ctx_lh_Lat_Fis-post | Cortical | 0.6012 | 0.0842 | 0.6170 | 0.0739 | 0.3562 | 0.0774 | 0.3474 | 0.0706 | 0.0426 | 0.0174 | 0.0355 | 0.0151 | 0.8373 | 0.0868 | 0.8525 | 0.0890 | 0.3724 | 0.0252 | 0.3803 | 0.0247 | 0.2531 | 0.0322 | 0.2696 | 0.0354 |
| ctx_lh_Pole_occipital | Cortical | 0.3210 | 0.1498 | 0.3034 | 0.1283 | 0.2007 | 0.1339 | 0.1952 | 0.1310 | 0.0602 | 0.0406 | 0.0500 | 0.0323 | 0.0488 | 0.0781 | 0.0467 | 0.0433 | 0.0001 | 0.0004 | 0.0002 | 0.0004 | 0.0000 | 0.0001 | 0.0001 | 0.0001 |
| ctx_lh_Pole_temporal | Cortical | 0.3593 | 0.1288 | 0.3597 | 0.1021 | 0.2002 | 0.0816 | 0.2205 | 0.0710 | 0.0416 | 0.0194 | 0.0440 | 0.0244 | 0.1926 | 0.0628 | 0.1918 | 0.0497 | 0.0370 | 0.0065 | 0.0388 | 0.0055 | 0.0247 | 0.0043 | 0.0258 | 0.0034 |
| ctx_lh_S_calcarine | Cortical | 0.5540 | 0.0586 | 0.5574 | 0.0621 | 0.3780 | 0.0594 | 0.3844 | 0.0630 | 0.0674 | 0.0262 | 0.0581 | 0.0196 | 0.8934 | 0.0908 | 0.9135 | 0.0950 | 0.4349 | 0.0258 | 0.4402 | 0.0255 | 0.2948 | 0.0301 | 0.3093 | 0.0348 |
| ctx_lh_S_central | Cortical | 0.5691 | 0.0465 | 0.5754 | 0.0525 | 0.3597 | 0.0434 | 0.3484 | 0.0549 | 0.0641 | 0.0264 | 0.0583 | 0.0286 | 0.7299 | 0.0909 | 0.7474 | 0.0929 | 0.3766 | 0.0254 | 0.3844 | 0.0244 | 0.2190 | 0.0254 | 0.2337 | 0.0263 |
| ctx_lh_S_cingul-Marginalis | Cortical | 0.5583 | 0.0656 | 0.5749 | 0.0687 | 0.3937 | 0.0582 | 0.3758 | 0.0637 | 0.0472 | 0.0251 | 0.0433 | 0.0251 | 0.8810 | 0.1032 | 0.8979 | 0.1029 | 0.4822 | 0.0307 | 0.4864 | 0.0309 | 0.3444 | 0.0374 | 0.3612 | 0.0441 |
| ctx_lh_S_circular_insula_ant | Cortical | 0.6607 | 0.0611 | 0.6602 | 0.0600 | 0.3051 | 0.0567 | 0.3112 | 0.0586 | 0.0342 | 0.0138 | 0.0283 | 0.0102 | 0.9294 | 0.1160 | 0.9486 | 0.1085 | 0.5224 | 0.0377 | 0.5275 | 0.0362 | 0.3427 | 0.0399 | 0.3586 | 0.0456 |
| ctx_lh_S_circular_insula_inf | Cortical | 0.5607 | 0.0636 | 0.5702 | 0.0645 | 0.3567 | 0.0522 | 0.3579 | 0.0573 | 0.0822 | 0.0364 | 0.0716 | 0.0290 | 0.8257 | 0.0867 | 0.8455 | 0.0874 | 0.4327 | 0.0314 | 0.4400 | 0.0268 | 0.2600 | 0.0293 | 0.2757 | 0.0337 |
| ctx_lh_S_circular_insula_sup | Cortical | 0.6069 | 0.0670 | 0.6102 | 0.0566 | 0.3583 | 0.0617 | 0.3588 | 0.0543 | 0.0348 | 0.0119 | 0.0309 | 0.0108 | 0.8709 | 0.0927 | 0.8882 | 0.0937 | 0.4749 | 0.0298 | 0.4818 | 0.0276 | 0.3112 | 0.0278 | 0.3245 | 0.0328 |
| ctx_lh_S_collat_transv_ant | Cortical | 0.6636 | 0.0750 | 0.6591 | 0.0826 | 0.2894 | 0.0691 | 0.3018 | 0.0815 | 0.0339 | 0.0101 | 0.0302 | 0.0100 | 0.9169 | 0.2127 | 0.9051 | 0.1565 | 0.2117 | 0.0239 | 0.2196 | 0.0218 | 0.1177 | 0.0150 | 0.1248 | 0.0177 |
| ctx_lh_S_collat_transv_post | Cortical | 0.5794 | 0.1106 | 0.5604 | 0.1102 | 0.3874 | 0.1005 | 0.4084 | 0.1039 | 0.0245 | 0.0240 | 0.0250 | 0.0221 | 1.1858 | 0.8967 | 1.2221 | 0.7122 | 0.1099 | 0.0331 | 0.1154 | 0.0334 | 0.0486 | 0.0130 | 0.0501 | 0.0140 |
| ctx_lh_S_front_inf | Cortical | 0.5812 | 0.0606 | 0.5870 | 0.0592 | 0.3778 | 0.0545 | 0.3760 | 0.0588 | 0.0409 | 0.0179 | 0.0369 | 0.0147 | 0.7472 | 0.1039 | 0.7689 | 0.0936 | 0.4146 | 0.0362 | 0.4164 | 0.0335 | 0.2096 | 0.0304 | 0.2180 | 0.0295 |
| ctx_lh_S_front_middle | Cortical | 0.5320 | 0.0550 | 0.5351 | 0.0514 | 0.4109 | 0.0484 | 0.4119 | 0.0503 | 0.0570 | 0.0304 | 0.0522 | 0.0245 | 0.6989 | 0.1156 | 0.7127 | 0.1078 | 0.4627 | 0.0554 | 0.4629 | 0.0509 | 0.2120 | 0.0337 | 0.2184 | 0.0355 |
| ctx_lh_S_front_sup | Cortical | 0.5632 | 0.0523 | 0.5616 | 0.0745 | 0.3895 | 0.0468 | 0.3796 | 0.0636 | 0.0469 | 0.0220 | 0.0421 | 0.0176 | 0.7573 | 0.1857 | 0.8006 | 0.1763 | 0.3143 | 0.0331 | 0.3171 | 0.0373 | 0.1555 | 0.0222 | 0.1630 | 0.0241 |
| ctx_lh_S_interm_prim-Jensen | Cortical | 0.5242 | 0.1041 | 0.5215 | 0.1017 | 0.4332 | 0.0940 | 0.4421 | 0.0950 | 0.0425 | 0.0284 | 0.0363 | 0.0232 | 0.7501 | 0.1566 | 0.7581 | 0.1404 | 0.4149 | 0.0682 | 0.4122 | 0.0744 | 0.1930 | 0.0416 | 0.1964 | 0.0481 |
| ctx_lh_S_intrapariet_and_P_trans | Cortical | 0.5199 | 0.0508 | 0.5336 | 0.0577 | 0.4155 | 0.0480 | 0.4035 | 0.0530 | 0.0640 | 0.0328 | 0.0623 | 0.0345 | 0.7567 | 0.0818 | 0.7763 | 0.0884 | 0.4051 | 0.0315 | 0.4130 | 0.0296 | 0.2272 | 0.0299 | 0.2429 | 0.0332 |
| ctx_lh_S_oc-temp_lat | Cortical | 0.5759 | 0.0743 | 0.5725 | 0.0628 | 0.3957 | 0.0734 | 0.4021 | 0.0626 | 0.0258 | 0.0114 | 0.0253 | 0.0105 | 0.7850 | 0.2195 | 0.8322 | 0.2003 | 0.3911 | 0.0340 | 0.3793 | 0.0323 | 0.1875 | 0.0250 | 0.1845 | 0.0244 |
| ctx_lh_S_oc-temp_med_and_Lingual | Cortical | 0.5935 | 0.0689 | 0.5929 | 0.0677 | 0.3789 | 0.0662 | 0.3843 | 0.0672 | 0.0273 | 0.0121 | 0.0228 | 0.0070 | 0.7964 | 0.0921 | 0.8155 | 0.0922 | 0.3094 | 0.0248 | 0.3114 | 0.0209 | 0.1566 | 0.0177 | 0.1613 | 0.0193 |
| ctx_lh_S_oc_middle_and_Lunatus | Cortical | 0.4835 | 0.0830 | 0.4911 | 0.0773 | 0.4310 | 0.0911 | 0.4406 | 0.0834 | 0.0545 | 0.0327 | 0.0501 | 0.0336 | 0.9123 | 0.5359 | 0.9711 | 0.4815 | 0.1885 | 0.0410 | 0.1926 | 0.0403 | 0.0839 | 0.0208 | 0.0896 | 0.0197 |
| ctx_lh_S_oc_sup_and_transversal | Cortical | 0.4951 | 0.0638 | 0.4944 | 0.0526 | 0.4283 | 0.0629 | 0.4392 | 0.0521 | 0.0652 | 0.0308 | 0.0613 | 0.0287 | 0.8588 | 0.1901 | 0.8772 | 0.1423 | 0.3861 | 0.0380 | 0.3925 | 0.0322 | 0.2098 | 0.0285 | 0.2217 | 0.0341 |
| ctx_lh_S_occipital_ant | Cortical | 0.5643 | 0.0689 | 0.5724 | 0.0713 | 0.4101 | 0.0656 | 0.4045 | 0.0697 | 0.0256 | 0.0103 | 0.0231 | 0.0105 | 0.8934 | 0.2466 | 0.8996 | 0.1710 | 0.3709 | 0.0366 | 0.3786 | 0.0390 | 0.1969 | 0.0288 | 0.2067 | 0.0318 |
| ctx_lh_S_orbital-H_Shaped | Cortical | 0.4935 | 0.1209 | 0.4520 | 0.1153 | 0.2563 | 0.0962 | 0.2616 | 0.0990 | 0.0739 | 0.0265 | 0.0689 | 0.0235 | 0.8371 | 0.1541 | 0.8237 | 0.1270 | 0.1636 | 0.0186 | 0.1649 | 0.0146 | 0.0891 | 0.0114 | 0.0909 | 0.0107 |
| ctx_lh_S_orbital_lateral | Cortical | 0.4917 | 0.0680 | 0.4922 | 0.0723 | 0.4192 | 0.0700 | 0.4254 | 0.0646 | 0.0675 | 0.0305 | 0.0617 | 0.0333 | 0.7029 | 0.2346 | 0.7596 | 0.2221 | 0.3437 | 0.0495 | 0.3470 | 0.0495 | 0.1451 | 0.0267 | 0.1554 | 0.0265 |
| ctx_lh_S_orbital_med-olfact | Cortical | 0.6041 | 0.1336 | 0.5926 | 0.1065 | 0.2097 | 0.0932 | 0.2255 | 0.0860 | 0.0396 | 0.0219 | 0.0404 | 0.0222 | 1.1732 | 0.3634 | 1.1163 | 0.2912 | 0.1948 | 0.0232 | 0.1963 | 0.0200 | 0.1335 | 0.0254 | 0.1363 | 0.0210 |
| ctx_lh_S_parieto_occipital | Cortical | 0.5875 | 0.0506 | 0.5932 | 0.0576 | 0.3627 | 0.0476 | 0.3601 | 0.0532 | 0.0497 | 0.0173 | 0.0467 | 0.0177 | 0.8886 | 0.0971 | 0.9030 | 0.0976 | 0.4398 | 0.0278 | 0.4435 | 0.0288 | 0.3094 | 0.0350 | 0.3218 | 0.0404 |
| ctx_lh_S_pericallosal | Cortical | 0.5843 | 0.0579 | 0.5867 | 0.0626 | 0.3734 | 0.0584 | 0.3758 | 0.0597 | 0.0423 | 0.0217 | 0.0373 | 0.0173 | 0.8897 | 0.1058 | 0.9168 | 0.1114 | 0.4158 | 0.0294 | 0.4203 | 0.0277 | 0.2546 | 0.0250 | 0.2663 | 0.0282 |
| ctx_lh_S_postcentral | Cortical | 0.5301 | 0.0543 | 0.5449 | 0.0479 | 0.4090 | 0.0482 | 0.3947 | 0.0505 | 0.0554 | 0.0262 | 0.0495 | 0.0263 | 0.7453 | 0.0846 | 0.7644 | 0.0873 | 0.4200 | 0.0410 | 0.4304 | 0.0406 | 0.2336 | 0.0322 | 0.2503 | 0.0383 |
| ctx_lh_S_precentral-inf-part | Cortical | 0.5706 | 0.0619 | 0.5663 | 0.0551 | 0.3883 | 0.0566 | 0.3949 | 0.0552 | 0.0410 | 0.0142 | 0.0383 | 0.0130 | 0.7161 | 0.0795 | 0.7242 | 0.0739 | 0.3706 | 0.0343 | 0.3750 | 0.0283 | 0.1872 | 0.0257 | 0.1937 | 0.0236 |
| ctx_lh_S_precentral-sup-part | Cortical | 0.5378 | 0.0615 | 0.5349 | 0.0898 | 0.4046 | 0.0544 | 0.3856 | 0.0824 | 0.0476 | 0.0226 | 0.0432 | 0.0224 | 0.6602 | 0.1330 | 0.7008 | 0.1365 | 0.2915 | 0.0291 | 0.2986 | 0.0297 | 0.1493 | 0.0190 | 0.1591 | 0.0186 |
| ctx_lh_S_suborbital | Cortical | 0.6106 | 0.1074 | 0.6061 | 0.0908 | 0.3018 | 0.0935 | 0.3186 | 0.0825 | 0.0706 | 0.0363 | 0.0632 | 0.0275 | 0.9843 | 0.1757 | 1.0118 | 0.1896 | 0.4708 | 0.0364 | 0.4713 | 0.0369 | 0.3154 | 0.0393 | 0.3266 | 0.0452 |
| ctx_lh_S_subparietal | Cortical | 0.5880 | 0.0669 | 0.5944 | 0.0644 | 0.3834 | 0.0626 | 0.3802 | 0.0607 | 0.0287 | 0.0128 | 0.0253 | 0.0125 | 0.9610 | 0.1132 | 0.9719 | 0.1124 | 0.4997 | 0.0354 | 0.5122 | 0.0337 | 0.3536 | 0.0410 | 0.3744 | 0.0442 |
| ctx_lh_S_temporal_inf | Cortical | 0.5851 | 0.0763 | 0.5785 | 0.0769 | 0.3809 | 0.0764 | 0.3943 | 0.0757 | 0.0296 | 0.0103 | 0.0263 | 0.0096 | 0.8486 | 0.3372 | 0.8530 | 0.2634 | 0.1620 | 0.0190 | 0.1599 | 0.0179 | 0.0914 | 0.0148 | 0.0901 | 0.0145 |
| ctx_lh_S_temporal_sup | Cortical | 0.5364 | 0.0570 | 0.5404 | 0.0576 | 0.4237 | 0.0545 | 0.4226 | 0.0585 | 0.0398 | 0.0129 | 0.0369 | 0.0134 | 0.8028 | 0.1334 | 0.8186 | 0.1029 | 0.3529 | 0.0258 | 0.3550 | 0.0229 | 0.1950 | 0.0208 | 0.2020 | 0.0205 |
| ctx_lh_S_temporal_transverse | Cortical | 0.5489 | 0.1039 | 0.5745 | 0.1090 | 0.3934 | 0.0851 | 0.3778 | 0.0986 | 0.0576 | 0.0364 | 0.0476 | 0.0315 | 1.0619 | 0.1289 | 1.0948 | 0.1281 | 0.5747 | 0.0392 | 0.5848 | 0.0357 | 0.4406 | 0.0603 | 0.4698 | 0.0604 |
| ctx_lh_Unknown | Cortical | 0.5632 | 0.0624 | 0.5672 | 0.0518 | 0.3289 | 0.0541 | 0.3372 | 0.0475 | 0.1052 | 0.0378 | 0.0950 | 0.0291 | 0.9153 | 0.2139 | 0.9234 | 0.1994 | 0.4465 | 0.0425 | 0.4491 | 0.0326 | 0.2587 | 0.0339 | 0.2648 | 0.0318 |
| ctx_rh_G_Ins_lg_and_S_cent_ins | Cortical | 0.4342 | 0.0546 | 0.4287 | 0.0490 | 0.4078 | 0.0531 | 0.4185 | 0.0527 | 0.1572 | 0.0465 | 0.1518 | 0.0402 | 0.5624 | 0.0803 | 0.5750 | 0.0750 | 0.1448 | 0.0392 | 0.1459 | 0.0238 | 0.0649 | 0.0149 | 0.0668 | 0.0109 |
| ctx_rh_G_and_S_cingul-Ant | Cortical | 0.6108 | 0.0532 | 0.6119 | 0.0422 | 0.3427 | 0.0506 | 0.3463 | 0.0422 | 0.0449 | 0.0134 | 0.0403 | 0.0124 | 0.9100 | 0.1026 | 0.9203 | 0.0982 | 0.4395 | 0.0276 | 0.4475 | 0.0230 | 0.3229 | 0.0332 | 0.3379 | 0.0380 |
| ctx_rh_G_and_S_cingul-Mid-Ant | Cortical | 0.5958 | 0.0573 | 0.5978 | 0.0638 | 0.3707 | 0.0518 | 0.3648 | 0.0533 | 0.0334 | 0.0125 | 0.0293 | 0.0125 | 0.8717 | 0.0998 | 0.8828 | 0.0987 | 0.5021 | 0.0324 | 0.5113 | 0.0264 | 0.3523 | 0.0354 | 0.3683 | 0.0376 |
| ctx_rh_G_and_S_cingul-Mid-Post | Cortical | 0.6065 | 0.0615 | 0.6169 | 0.0720 | 0.3623 | 0.0566 | 0.3502 | 0.0618 | 0.0312 | 0.0122 | 0.0265 | 0.0129 | 0.9025 | 0.0987 | 0.9104 | 0.0965 | 0.5030 | 0.0326 | 0.5165 | 0.0306 | 0.3658 | 0.0395 | 0.3864 | 0.0430 |
| ctx_rh_G_and_S_frontomargin | Cortical | 0.3781 | 0.1556 | 0.3672 | 0.1652 | 0.1578 | 0.0822 | 0.1621 | 0.0828 | 0.0609 | 0.0383 | 0.0490 | 0.0267 | 1.0302 | 0.5176 | 1.0225 | 0.4602 | 0.1709 | 0.0257 | 0.1721 | 0.0228 | 0.0770 | 0.0119 | 0.0795 | 0.0117 |
| ctx_rh_G_and_S_occipital_inf | Cortical | 0.5698 | 0.0967 | 0.5673 | 0.1108 | 0.2752 | 0.0804 | 0.2638 | 0.0874 | 0.0638 | 0.0327 | 0.0668 | 0.0425 | 0.6003 | 0.3747 | 0.6249 | 0.3449 | 0.0499 | 0.0125 | 0.0495 | 0.0134 | 0.0257 | 0.0070 | 0.0260 | 0.0068 |
| ctx_rh_G_and_S_paracentral | Cortical | 0.3545 | 0.1028 | 0.3449 | 0.1108 | 0.2258 | 0.0614 | 0.2161 | 0.0710 | 0.1395 | 0.0822 | 0.1294 | 0.0870 | 0.3657 | 0.0848 | 0.3916 | 0.0992 | 0.1581 | 0.0122 | 0.1590 | 0.0112 | 0.0986 | 0.0120 | 0.1023 | 0.0126 |
| ctx_rh_G_and_S_subcentral | Cortical | 0.5133 | 0.0695 | 0.5120 | 0.0547 | 0.4018 | 0.0593 | 0.4134 | 0.0489 | 0.0834 | 0.0385 | 0.0741 | 0.0285 | 0.7634 | 0.1495 | 0.7887 | 0.1388 | 0.2232 | 0.0175 | 0.2280 | 0.0169 | 0.1397 | 0.0158 | 0.1479 | 0.0183 |
| ctx_rh_G_and_S_transv_frontopol | Cortical | 0.4514 | 0.1335 | 0.4279 | 0.1496 | 0.2061 | 0.0785 | 0.2091 | 0.0851 | 0.0914 | 0.0446 | 0.0833 | 0.0414 | 0.8119 | 0.4581 | 0.8673 | 0.4931 | 0.2045 | 0.0316 | 0.2051 | 0.0298 | 0.0873 | 0.0139 | 0.0904 | 0.0133 |
| ctx_rh_G_cingul-Post-dorsal | Cortical | 0.5660 | 0.0794 | 0.5761 | 0.0696 | 0.4125 | 0.0757 | 0.4055 | 0.0693 | 0.0216 | 0.0116 | 0.0181 | 0.0103 | 0.7611 | 0.0788 | 0.7647 | 0.0872 | 0.3793 | 0.0342 | 0.3880 | 0.0335 | 0.1778 | 0.0200 | 0.1853 | 0.0216 |
| ctx_rh_G_cingul-Post-ventral | Cortical | 0.5819 | 0.1035 | 0.5846 | 0.0948 | 0.3674 | 0.0921 | 0.3702 | 0.0903 | 0.0506 | 0.0399 | 0.0452 | 0.0276 | 0.7522 | 0.1172 | 0.7717 | 0.1076 | 0.1859 | 0.0245 | 0.1880 | 0.0208 | 0.0858 | 0.0130 | 0.0874 | 0.0121 |
| ctx_rh_G_cuneus | Cortical | 0.5300 | 0.0576 | 0.5359 | 0.0647 | 0.3773 | 0.0528 | 0.3802 | 0.0611 | 0.0869 | 0.0307 | 0.0803 | 0.0331 | 0.9388 | 0.1454 | 0.9659 | 0.1518 | 0.3158 | 0.0267 | 0.3192 | 0.0259 | 0.1880 | 0.0260 | 0.1975 | 0.0297 |
| ctx_rh_G_front_inf-Opercular | Cortical | 0.4828 | 0.0488 | 0.4850 | 0.0442 | 0.4203 | 0.0485 | 0.4299 | 0.0461 | 0.0963 | 0.0402 | 0.0845 | 0.0309 | 0.8733 | 0.2059 | 0.8937 | 0.1922 | 0.2325 | 0.0214 | 0.2377 | 0.0224 | 0.1193 | 0.0136 | 0.1253 | 0.0147 |
| ctx_rh_G_front_inf-Orbital | Cortical | 0.5327 | 0.0929 | 0.5243 | 0.0918 | 0.3409 | 0.0763 | 0.3529 | 0.0739 | 0.0641 | 0.0303 | 0.0593 | 0.0296 | 0.6792 | 0.1456 | 0.7039 | 0.1465 | 0.2815 | 0.0324 | 0.2850 | 0.0281 | 0.1668 | 0.0303 | 0.1772 | 0.0315 |
| ctx_rh_G_front_inf-Triangul | Cortical | 0.5154 | 0.0493 | 0.5145 | 0.0452 | 0.4042 | 0.0495 | 0.4136 | 0.0453 | 0.0753 | 0.0310 | 0.0673 | 0.0271 | 0.8776 | 0.3323 | 0.9292 | 0.3163 | 0.2349 | 0.0294 | 0.2320 | 0.0280 | 0.1181 | 0.0169 | 0.1210 | 0.0173 |
| ctx_rh_G_front_middle | Cortical | 0.4996 | 0.0466 | 0.4937 | 0.0538 | 0.4058 | 0.0513 | 0.4101 | 0.0551 | 0.0863 | 0.0345 | 0.0799 | 0.0288 | 0.7887 | 0.4267 | 0.8466 | 0.4133 | 0.1552 | 0.0207 | 0.1557 | 0.0208 | 0.0717 | 0.0107 | 0.0748 | 0.0099 |
| ctx_rh_G_front_sup | Cortical | 0.4883 | 0.0562 | 0.4757 | 0.0723 | 0.3200 | 0.0475 | 0.3145 | 0.0623 | 0.1202 | 0.0401 | 0.1096 | 0.0413 | 0.5618 | 0.2146 | 0.6057 | 0.2055 | 0.2176 | 0.0147 | 0.2190 | 0.0133 | 0.1228 | 0.0120 | 0.1292 | 0.0133 |
| ctx_rh_G_insular_short | Cortical | 0.4631 | 0.0487 | 0.4630 | 0.0434 | 0.4155 | 0.0471 | 0.4294 | 0.0449 | 0.1209 | 0.0454 | 0.1070 | 0.0353 | 0.6219 | 0.1039 | 0.6396 | 0.0965 | 0.1306 | 0.0287 | 0.1348 | 0.0252 | 0.0537 | 0.0117 | 0.0565 | 0.0105 |
| ctx_rh_G_oc-temp_lat-fusifor | Cortical | 0.5465 | 0.0766 | 0.5331 | 0.0803 | 0.3703 | 0.0653 | 0.3779 | 0.0702 | 0.0737 | 0.0278 | 0.0794 | 0.0272 | 0.7804 | 0.1388 | 0.7958 | 0.1198 | 0.4064 | 0.0457 | 0.4128 | 0.0356 | 0.1978 | 0.0263 | 0.2049 | 0.0247 |
| ctx_rh_G_oc-temp_med-Lingual | Cortical | 0.5543 | 0.0724 | 0.5616 | 0.0755 | 0.3791 | 0.0643 | 0.3774 | 0.0712 | 0.0626 | 0.0261 | 0.0578 | 0.0243 | 0.8355 | 0.1227 | 0.8539 | 0.1248 | 0.2492 | 0.0213 | 0.2527 | 0.0213 | 0.1277 | 0.0139 | 0.1340 | 0.0151 |
| ctx_rh_G_oc-temp_med-Parahip | Cortical | 0.5501 | 0.0704 | 0.5441 | 0.0734 | 0.3243 | 0.0679 | 0.3467 | 0.0620 | 0.0987 | 0.0316 | 0.0954 | 0.0355 | 1.2721 | 0.2963 | 1.2960 | 0.2876 | 0.2565 | 0.0248 | 0.2587 | 0.0224 | 0.1702 | 0.0198 | 0.1737 | 0.0218 |
| ctx_rh_G_occipital_middle | Cortical | 0.5066 | 0.0880 | 0.5070 | 0.0824 | 0.2517 | 0.0932 | 0.2560 | 0.0933 | 0.0856 | 0.0376 | 0.0838 | 0.0356 | 0.7588 | 0.4774 | 0.7978 | 0.4107 | 0.0559 | 0.0139 | 0.0565 | 0.0138 | 0.0267 | 0.0066 | 0.0290 | 0.0069 |
| ctx_rh_G_occipital_sup | Cortical | 0.4634 | 0.1121 | 0.4552 | 0.1000 | 0.2111 | 0.1017 | 0.2219 | 0.0976 | 0.1407 | 0.0568 | 0.1378 | 0.0624 | 0.8593 | 0.4424 | 0.9261 | 0.4189 | 0.1615 | 0.0224 | 0.1631 | 0.0233 | 0.0789 | 0.0138 | 0.0835 | 0.0133 |
| ctx_rh_G_orbital | Cortical | 0.4131 | 0.1123 | 0.3939 | 0.1209 | 0.1866 | 0.0756 | 0.1900 | 0.0755 | 0.0735 | 0.0231 | 0.0692 | 0.0222 | 0.7701 | 0.1781 | 0.8097 | 0.1857 | 0.0968 | 0.0112 | 0.0996 | 0.0116 | 0.0503 | 0.0056 | 0.0529 | 0.0067 |
| ctx_rh_G_pariet_inf-Angular | Cortical | 0.4804 | 0.0680 | 0.4776 | 0.0586 | 0.3750 | 0.0790 | 0.3866 | 0.0727 | 0.1088 | 0.0370 | 0.1029 | 0.0344 | 0.8043 | 0.3755 | 0.8429 | 0.3621 | 0.1798 | 0.0260 | 0.1828 | 0.0258 | 0.0848 | 0.0155 | 0.0892 | 0.0134 |
| ctx_rh_G_pariet_inf-Supramar | Cortical | 0.4649 | 0.0500 | 0.4711 | 0.0457 | 0.4574 | 0.0485 | 0.4600 | 0.0497 | 0.0760 | 0.0318 | 0.0666 | 0.0233 | 0.7473 | 0.2505 | 0.7727 | 0.2381 | 0.1906 | 0.0212 | 0.1958 | 0.0203 | 0.0902 | 0.0132 | 0.0960 | 0.0131 |
| ctx_rh_G_parietal_sup | Cortical | 0.4008 | 0.0953 | 0.3902 | 0.0917 | 0.2881 | 0.0885 | 0.2920 | 0.0842 | 0.1813 | 0.0723 | 0.1790 | 0.0719 | 0.4552 | 0.1934 | 0.4519 | 0.1818 | 0.1365 | 0.0181 | 0.1395 | 0.0169 | 0.0613 | 0.0103 | 0.0642 | 0.0097 |
| ctx_rh_G_postcentral | Cortical | 0.3791 | 0.0664 | 0.3707 | 0.0685 | 0.3404 | 0.0698 | 0.3428 | 0.0676 | 0.1735 | 0.0645 | 0.1657 | 0.0631 | 0.3342 | 0.1684 | 0.3464 | 0.1494 | 0.0341 | 0.0067 | 0.0353 | 0.0070 | 0.0157 | 0.0029 | 0.0172 | 0.0031 |
| ctx_rh_G_precentral | Cortical | 0.4099 | 0.0599 | 0.3994 | 0.0688 | 0.3463 | 0.0563 | 0.3463 | 0.0639 | 0.1491 | 0.0520 | 0.1355 | 0.0501 | 0.4215 | 0.1913 | 0.4451 | 0.1691 | 0.0625 | 0.0097 | 0.0645 | 0.0092 | 0.0299 | 0.0052 | 0.0321 | 0.0049 |
| ctx_rh_G_precuneus | Cortical | 0.4707 | 0.0546 | 0.4712 | 0.0539 | 0.3720 | 0.0528 | 0.3743 | 0.0609 | 0.1261 | 0.0482 | 0.1179 | 0.0451 | 0.7180 | 0.1079 | 0.7363 | 0.1039 | 0.2881 | 0.0232 | 0.2905 | 0.0210 | 0.1459 | 0.0161 | 0.1533 | 0.0173 |
| ctx_rh_G_rectus | Cortical | 0.4232 | 0.1334 | 0.4072 | 0.1338 | 0.1927 | 0.0947 | 0.2012 | 0.0984 | 0.1074 | 0.0532 | 0.1088 | 0.0464 | 1.2008 | 0.4704 | 1.2780 | 0.5066 | 0.0789 | 0.0136 | 0.0789 | 0.0125 | 0.0502 | 0.0093 | 0.0530 | 0.0105 |
| ctx_rh_G_subcallosal | Cortical | 0.5358 | 0.0692 | 0.5339 | 0.0585 | 0.3899 | 0.0702 | 0.3979 | 0.0591 | 0.0665 | 0.0341 | 0.0602 | 0.0297 | 0.9307 | 0.2308 | 0.9109 | 0.1688 | 0.2721 | 0.0431 | 0.2866 | 0.0400 | 0.1579 | 0.0403 | 0.1716 | 0.0473 |
| ctx_rh_G_temp_sup-G_T_transv | Cortical | 0.4391 | 0.0620 | 0.4356 | 0.0728 | 0.4200 | 0.0555 | 0.4376 | 0.0624 | 0.1404 | 0.0521 | 0.1262 | 0.0491 | 0.7498 | 0.0837 | 0.7722 | 0.0812 | 0.2797 | 0.0214 | 0.2829 | 0.0198 | 0.1493 | 0.0160 | 0.1565 | 0.0188 |
| ctx_rh_G_temp_sup-Lateral | Cortical | 0.4883 | 0.0554 | 0.4809 | 0.0480 | 0.4032 | 0.0555 | 0.4162 | 0.0550 | 0.0933 | 0.0349 | 0.0870 | 0.0315 | 0.6634 | 0.2360 | 0.7005 | 0.2367 | 0.0854 | 0.0096 | 0.0860 | 0.0095 | 0.0423 | 0.0057 | 0.0435 | 0.0060 |
| ctx_rh_G_temp_sup-Plan_polar | Cortical | 0.5416 | 0.0797 | 0.5442 | 0.0746 | 0.3336 | 0.0563 | 0.3479 | 0.0536 | 0.1175 | 0.0565 | 0.1026 | 0.0505 | 0.9006 | 0.1268 | 0.9279 | 0.1211 | 0.5285 | 0.0393 | 0.5309 | 0.0358 | 0.3463 | 0.0354 | 0.3588 | 0.0397 |
| ctx_rh_G_temp_sup-Plan_tempo | Cortical | 0.5206 | 0.0679 | 0.5350 | 0.0710 | 0.4247 | 0.0591 | 0.4182 | 0.0644 | 0.0546 | 0.0291 | 0.0468 | 0.0255 | 0.8935 | 0.1623 | 0.9175 | 0.1620 | 0.4417 | 0.0329 | 0.4477 | 0.0304 | 0.2545 | 0.0296 | 0.2666 | 0.0335 |
| ctx_rh_G_temporal_inf | Cortical | 0.5537 | 0.0866 | 0.5502 | 0.0843 | 0.2992 | 0.0704 | 0.3157 | 0.0741 | 0.0529 | 0.0169 | 0.0543 | 0.0186 | 0.7280 | 0.2767 | 0.7568 | 0.2653 | 0.0756 | 0.0124 | 0.0757 | 0.0126 | 0.0423 | 0.0083 | 0.0422 | 0.0057 |
| ctx_rh_G_temporal_middle | Cortical | 0.5160 | 0.0652 | 0.5099 | 0.0572 | 0.4110 | 0.0635 | 0.4255 | 0.0555 | 0.0548 | 0.0185 | 0.0505 | 0.0133 | 0.5788 | 0.2290 | 0.5968 | 0.2132 | 0.0907 | 0.0113 | 0.0923 | 0.0113 | 0.0460 | 0.0071 | 0.0475 | 0.0072 |
| ctx_rh_Lat_Fis-ant-Horizont | Cortical | 0.7125 | 0.0863 | 0.7291 | 0.0828 | 0.2515 | 0.0796 | 0.2391 | 0.0743 | 0.0351 | 0.0158 | 0.0314 | 0.0164 | 0.9935 | 0.1229 | 1.0193 | 0.1273 | 0.6335 | 0.0336 | 0.6413 | 0.0373 | 0.4905 | 0.0543 | 0.5156 | 0.0680 |
| ctx_rh_Lat_Fis-ant-Vertical | Cortical | 0.6390 | 0.1296 | 0.6453 | 0.1187 | 0.3183 | 0.1146 | 0.3150 | 0.1042 | 0.0426 | 0.0288 | 0.0397 | 0.0267 | 1.0240 | 0.1264 | 1.0370 | 0.1361 | 0.6470 | 0.0362 | 0.6489 | 0.0401 | 0.5064 | 0.0652 | 0.5187 | 0.0741 |
| ctx_rh_Lat_Fis-post | Cortical | 0.5701 | 0.0760 | 0.5792 | 0.0706 | 0.3838 | 0.0686 | 0.3797 | 0.0661 | 0.0461 | 0.0198 | 0.0410 | 0.0170 | 0.7900 | 0.0842 | 0.8057 | 0.0828 | 0.3740 | 0.0264 | 0.3798 | 0.0245 | 0.2198 | 0.0240 | 0.2320 | 0.0261 |
| ctx_rh_Pole_occipital | Cortical | 0.4080 | 0.1364 | 0.3873 | 0.1198 | 0.1846 | 0.1022 | 0.1800 | 0.1032 | 0.0684 | 0.0376 | 0.0657 | 0.0368 | 0.1875 | 0.1188 | 0.1865 | 0.1075 | 0.0115 | 0.0042 | 0.0116 | 0.0038 | 0.0038 | 0.0015 | 0.0043 | 0.0016 |
| ctx_rh_Pole_temporal | Cortical | 0.3448 | 0.1623 | 0.3702 | 0.1161 | 0.2014 | 0.0987 | 0.2306 | 0.0785 | 0.0413 | 0.0213 | 0.0470 | 0.0272 | 0.2245 | 0.0895 | 0.2299 | 0.0778 | 0.0346 | 0.0071 | 0.0354 | 0.0061 | 0.0232 | 0.0051 | 0.0234 | 0.0037 |
| ctx_rh_S_calcarine | Cortical | 0.5813 | 0.0713 | 0.5901 | 0.0759 | 0.3531 | 0.0688 | 0.3578 | 0.0736 | 0.0652 | 0.0407 | 0.0521 | 0.0148 | 0.9262 | 0.0970 | 0.9459 | 0.1013 | 0.4323 | 0.0270 | 0.4370 | 0.0225 | 0.2917 | 0.0307 | 0.3056 | 0.0352 |
| ctx_rh_S_central | Cortical | 0.5514 | 0.0495 | 0.5563 | 0.0560 | 0.3748 | 0.0432 | 0.3684 | 0.0553 | 0.0686 | 0.0304 | 0.0603 | 0.0285 | 0.7191 | 0.0916 | 0.7420 | 0.0916 | 0.3520 | 0.0274 | 0.3617 | 0.0265 | 0.1847 | 0.0209 | 0.1993 | 0.0221 |
| ctx_rh_S_cingul-Marginalis | Cortical | 0.5944 | 0.0679 | 0.6007 | 0.0746 | 0.3589 | 0.0606 | 0.3501 | 0.0630 | 0.0448 | 0.0260 | 0.0404 | 0.0223 | 0.8645 | 0.0953 | 0.8808 | 0.1001 | 0.4687 | 0.0302 | 0.4771 | 0.0328 | 0.3262 | 0.0362 | 0.3455 | 0.0440 |
| ctx_rh_S_circular_insula_ant | Cortical | 0.6214 | 0.0593 | 0.6284 | 0.0540 | 0.3389 | 0.0556 | 0.3366 | 0.0519 | 0.0395 | 0.0143 | 0.0339 | 0.0116 | 0.9609 | 0.1197 | 0.9784 | 0.1146 | 0.4687 | 0.0347 | 0.4781 | 0.0298 | 0.3098 | 0.0368 | 0.3306 | 0.0428 |
| ctx_rh_S_circular_insula_inf | Cortical | 0.5555 | 0.0709 | 0.5547 | 0.0688 | 0.3669 | 0.0537 | 0.3736 | 0.0566 | 0.0773 | 0.0355 | 0.0708 | 0.0338 | 0.8449 | 0.0939 | 0.8734 | 0.0939 | 0.4274 | 0.0326 | 0.4318 | 0.0255 | 0.2669 | 0.0271 | 0.2825 | 0.0300 |
| ctx_rh_S_circular_insula_sup | Cortical | 0.6017 | 0.0692 | 0.5983 | 0.0552 | 0.3632 | 0.0648 | 0.3695 | 0.0523 | 0.0351 | 0.0116 | 0.0321 | 0.0117 | 0.8770 | 0.0954 | 0.8977 | 0.0956 | 0.4969 | 0.0302 | 0.5026 | 0.0269 | 0.3323 | 0.0298 | 0.3475 | 0.0345 |
| ctx_rh_S_collat_transv_ant | Cortical | 0.6232 | 0.0901 | 0.6258 | 0.0969 | 0.3153 | 0.0758 | 0.3220 | 0.0829 | 0.0350 | 0.0110 | 0.0323 | 0.0111 | 1.0691 | 0.2251 | 1.0739 | 0.1964 | 0.2591 | 0.0214 | 0.2623 | 0.0211 | 0.1550 | 0.0177 | 0.1612 | 0.0195 |
| ctx_rh_S_collat_transv_post | Cortical | 0.6462 | 0.1261 | 0.6359 | 0.1274 | 0.3199 | 0.1185 | 0.3331 | 0.1173 | 0.0286 | 0.0217 | 0.0258 | 0.0171 | 1.0515 | 0.6061 | 1.0974 | 0.5163 | 0.1983 | 0.0378 | 0.1962 | 0.0330 | 0.0902 | 0.0175 | 0.0920 | 0.0173 |
| ctx_rh_S_front_inf | Cortical | 0.6014 | 0.0608 | 0.6027 | 0.0613 | 0.3599 | 0.0582 | 0.3629 | 0.0615 | 0.0386 | 0.0143 | 0.0341 | 0.0137 | 0.7821 | 0.1127 | 0.8045 | 0.1033 | 0.4731 | 0.0389 | 0.4792 | 0.0384 | 0.2571 | 0.0346 | 0.2694 | 0.0365 |
| ctx_rh_S_front_middle | Cortical | 0.5498 | 0.0559 | 0.5476 | 0.0497 | 0.3992 | 0.0501 | 0.4049 | 0.0504 | 0.0507 | 0.0256 | 0.0457 | 0.0225 | 0.7330 | 0.1956 | 0.7614 | 0.2107 | 0.3989 | 0.0437 | 0.4008 | 0.0423 | 0.1807 | 0.0273 | 0.1873 | 0.0255 |
| ctx_rh_S_front_sup | Cortical | 0.5625 | 0.0523 | 0.5562 | 0.0812 | 0.3892 | 0.0477 | 0.3835 | 0.0681 | 0.0477 | 0.0200 | 0.0430 | 0.0208 | 0.7652 | 0.2517 | 0.8037 | 0.2720 | 0.2827 | 0.0343 | 0.2872 | 0.0321 | 0.1361 | 0.0196 | 0.1439 | 0.0201 |
| ctx_rh_S_interm_prim-Jensen | Cortical | 0.5384 | 0.0880 | 0.5536 | 0.0990 | 0.4173 | 0.0754 | 0.4048 | 0.0894 | 0.0444 | 0.0342 | 0.0385 | 0.0304 | 0.7374 | 0.0963 | 0.7519 | 0.0952 | 0.5153 | 0.0697 | 0.5141 | 0.0807 | 0.2630 | 0.0535 | 0.2736 | 0.0683 |
| ctx_rh_S_intrapariet_and_P_trans | Cortical | 0.5377 | 0.0533 | 0.5462 | 0.0597 | 0.4018 | 0.0485 | 0.3961 | 0.0585 | 0.0602 | 0.0324 | 0.0545 | 0.0301 | 0.7573 | 0.0865 | 0.7761 | 0.0884 | 0.4407 | 0.0400 | 0.4519 | 0.0362 | 0.2335 | 0.0306 | 0.2501 | 0.0346 |
| ctx_rh_S_oc-temp_lat | Cortical | 0.6032 | 0.0763 | 0.5988 | 0.0818 | 0.3738 | 0.0754 | 0.3791 | 0.0821 | 0.0217 | 0.0073 | 0.0199 | 0.0074 | 0.7952 | 0.1608 | 0.8173 | 0.1626 | 0.3559 | 0.0289 | 0.3518 | 0.0287 | 0.1865 | 0.0260 | 0.1850 | 0.0234 |
| ctx_rh_S_oc-temp_med_and_Lingual | Cortical | 0.6071 | 0.0794 | 0.6035 | 0.0774 | 0.3646 | 0.0762 | 0.3721 | 0.0763 | 0.0281 | 0.0146 | 0.0239 | 0.0079 | 0.8192 | 0.0926 | 0.8393 | 0.0909 | 0.3078 | 0.0264 | 0.3091 | 0.0247 | 0.1626 | 0.0192 | 0.1694 | 0.0222 |
| ctx_rh_S_oc_middle_and_Lunatus | Cortical | 0.5741 | 0.1148 | 0.5751 | 0.1198 | 0.3791 | 0.1063 | 0.3845 | 0.1136 | 0.0411 | 0.0290 | 0.0357 | 0.0214 | 0.9628 | 0.5101 | 0.9858 | 0.3566 | 0.3931 | 0.0436 | 0.3917 | 0.0428 | 0.2238 | 0.0366 | 0.2278 | 0.0361 |
| ctx_rh_S_oc_sup_and_transversal | Cortical | 0.5465 | 0.0681 | 0.5503 | 0.0847 | 0.3865 | 0.0632 | 0.3933 | 0.0784 | 0.0585 | 0.0271 | 0.0513 | 0.0242 | 0.8693 | 0.1555 | 0.8874 | 0.1289 | 0.4108 | 0.0334 | 0.4122 | 0.0309 | 0.2428 | 0.0330 | 0.2493 | 0.0339 |
| ctx_rh_S_occipital_ant | Cortical | 0.6149 | 0.1012 | 0.6261 | 0.1111 | 0.3625 | 0.0964 | 0.3547 | 0.1073 | 0.0226 | 0.0103 | 0.0193 | 0.0080 | 0.8809 | 0.2558 | 0.8860 | 0.1612 | 0.4397 | 0.0435 | 0.4420 | 0.0432 | 0.2499 | 0.0370 | 0.2572 | 0.0389 |
| ctx_rh_S_orbital-H_Shaped | Cortical | 0.5050 | 0.1151 | 0.4707 | 0.1132 | 0.2810 | 0.0970 | 0.2901 | 0.0919 | 0.0670 | 0.0249 | 0.0668 | 0.0248 | 0.9232 | 0.1574 | 0.9114 | 0.1438 | 0.2029 | 0.0196 | 0.2054 | 0.0181 | 0.1155 | 0.0142 | 0.1189 | 0.0155 |
| ctx_rh_S_orbital_lateral | Cortical | 0.5445 | 0.0838 | 0.5452 | 0.0834 | 0.3990 | 0.0753 | 0.4037 | 0.0760 | 0.0480 | 0.0230 | 0.0415 | 0.0217 | 0.7377 | 0.2185 | 0.7490 | 0.2066 | 0.3588 | 0.0479 | 0.3573 | 0.0522 | 0.1623 | 0.0307 | 0.1643 | 0.0343 |
| ctx_rh_S_orbital_med-olfact | Cortical | 0.5918 | 0.1131 | 0.5812 | 0.1006 | 0.2617 | 0.0883 | 0.2695 | 0.0810 | 0.0523 | 0.0280 | 0.0477 | 0.0286 | 1.0158 | 0.2503 | 1.0607 | 0.2491 | 0.2513 | 0.0264 | 0.2574 | 0.0259 | 0.1601 | 0.0231 | 0.1665 | 0.0218 |
| ctx_rh_S_parieto_occipital | Cortical | 0.6068 | 0.0592 | 0.6125 | 0.0728 | 0.3457 | 0.0543 | 0.3423 | 0.0686 | 0.0475 | 0.0174 | 0.0452 | 0.0178 | 0.8622 | 0.0932 | 0.8752 | 0.0964 | 0.3930 | 0.0276 | 0.3994 | 0.0249 | 0.2645 | 0.0328 | 0.2791 | 0.0372 |
| ctx_rh_S_pericallosal | Cortical | 0.5452 | 0.0499 | 0.5504 | 0.0521 | 0.4111 | 0.0495 | 0.4105 | 0.0496 | 0.0436 | 0.0193 | 0.0389 | 0.0142 | 0.8167 | 0.0921 | 0.8351 | 0.0916 | 0.4061 | 0.0311 | 0.4138 | 0.0287 | 0.2193 | 0.0225 | 0.2309 | 0.0246 |
| ctx_rh_S_postcentral | Cortical | 0.5162 | 0.0523 | 0.5278 | 0.0584 | 0.4195 | 0.0495 | 0.4067 | 0.0589 | 0.0601 | 0.0307 | 0.0536 | 0.0278 | 0.6972 | 0.0869 | 0.7158 | 0.0849 | 0.3881 | 0.0437 | 0.4011 | 0.0420 | 0.1950 | 0.0282 | 0.2116 | 0.0337 |
| ctx_rh_S_precentral-inf-part | Cortical | 0.5571 | 0.0596 | 0.5592 | 0.0593 | 0.3970 | 0.0545 | 0.3978 | 0.0583 | 0.0459 | 0.0182 | 0.0406 | 0.0165 | 0.7115 | 0.0867 | 0.7304 | 0.0873 | 0.3214 | 0.0340 | 0.3245 | 0.0342 | 0.1582 | 0.0254 | 0.1651 | 0.0247 |
| ctx_rh_S_precentral-sup-part | Cortical | 0.5384 | 0.0579 | 0.5413 | 0.0838 | 0.4032 | 0.0528 | 0.3849 | 0.0779 | 0.0489 | 0.0230 | 0.0428 | 0.0224 | 0.6851 | 0.1489 | 0.7052 | 0.1681 | 0.3232 | 0.0365 | 0.3325 | 0.0394 | 0.1631 | 0.0214 | 0.1742 | 0.0247 |
| ctx_rh_S_suborbital | Cortical | 0.5671 | 0.1166 | 0.5567 | 0.1119 | 0.3099 | 0.1125 | 0.3200 | 0.1107 | 0.0853 | 0.0542 | 0.0937 | 0.0610 | 1.4180 | 0.5194 | 1.5154 | 0.5090 | 0.3689 | 0.0333 | 0.3692 | 0.0318 | 0.2455 | 0.0324 | 0.2541 | 0.0332 |
| ctx_rh_S_subparietal | Cortical | 0.5779 | 0.0682 | 0.5847 | 0.0739 | 0.3932 | 0.0635 | 0.3879 | 0.0703 | 0.0289 | 0.0126 | 0.0269 | 0.0127 | 0.9061 | 0.1058 | 0.9142 | 0.1065 | 0.4414 | 0.0360 | 0.4524 | 0.0345 | 0.2699 | 0.0326 | 0.2864 | 0.0359 |
| ctx_rh_S_temporal_inf | Cortical | 0.5819 | 0.0825 | 0.5897 | 0.0814 | 0.3782 | 0.0798 | 0.3772 | 0.0773 | 0.0276 | 0.0094 | 0.0249 | 0.0081 | 0.9161 | 0.2639 | 0.9090 | 0.2566 | 0.2423 | 0.0222 | 0.2398 | 0.0267 | 0.1379 | 0.0190 | 0.1385 | 0.0228 |
| ctx_rh_S_temporal_sup | Cortical | 0.5350 | 0.0574 | 0.5420 | 0.0597 | 0.4272 | 0.0553 | 0.4242 | 0.0594 | 0.0375 | 0.0117 | 0.0335 | 0.0108 | 0.7968 | 0.0990 | 0.8153 | 0.0915 | 0.3431 | 0.0210 | 0.3449 | 0.0193 | 0.1989 | 0.0198 | 0.2071 | 0.0217 |
| ctx_rh_S_temporal_transverse | Cortical | 0.5862 | 0.1339 | 0.6253 | 0.1417 | 0.3574 | 0.1121 | 0.3302 | 0.1202 | 0.0563 | 0.0446 | 0.0445 | 0.0370 | 1.1366 | 0.1393 | 1.1759 | 0.1372 | 0.5901 | 0.0405 | 0.5960 | 0.0360 | 0.4711 | 0.0594 | 0.5003 | 0.0672 |
| ctx_rh_Unknown | Cortical | 0.5480 | 0.0650 | 0.5400 | 0.0655 | 0.3020 | 0.0612 | 0.3244 | 0.0619 | 0.1482 | 0.0545 | 0.1301 | 0.0468 | 1.0137 | 0.3937 | 1.0503 | 0.3322 | 0.3228 | 0.0424 | 0.3294 | 0.0422 | 0.1757 | 0.0288 | 0.1829 | 0.0322 |

*Supplementary Table 1: Summary means and standard deviations from each of the 214 ROIs in this study from each microstructural metric.*
